# Comparative population genomics reveals adaptive convergence in two *Drosophila* species across global environments

**DOI:** 10.1101/2025.10.02.680045

**Authors:** Weixuan Li, Chenlu Liu, Junhao Chen, Qinuo Wang, Yirong Wang, Andrew G. Clark, Jian Lu

## Abstract

The extent to which evolution is predictable across lineages remains unclear. To address this question, we investigated convergent adaptation in a pair of globally distributed sibling species, *Drosophila melanogaster* and *D. simulans*. We integrated whole-genome data from approximately 2,000 strains sampled across major continents, and revealed a more recent global colonization of *D. simulans* relative to *D. melanogaster*. Using a suite of complementary selection scans, we quantified signatures of positive selection across evolutionary timescales and genomic contexts. Despite substantial divergence, approximately 9–13% of adaptively evolving genes were shared between species across methods, revealing widespread convergence at the gene and pathway levels. Convergence was particularly pronounced for insecticide resistance genes, and was also evident in oxidative stress experiment. This study provides a quantitative, multiscale framework for dissecting molecular convergence, offering insights into the predictability of evolution, the constraints imposed by genomic architecture, and the dynamics of adaptation under global environmental change.

## Introduction

The extent to which evolution is repeatable is a fundamental question that continues to be debated in the field of evolutionary biology^1^. While natural selection is expected to drive similar outcomes in similar environments^2,3^, the unpredictability of mutation, recombination, demographic history, and historical contingency often leads to divergent evolutionary trajectories^4^. This tension between determinism and chance shapes the course of evolution and underpins the broader question of whether adaptive outcomes are predictable.

Studies of evolutionary convergence offer a powerful lens into this issue. Experimental evolution has revealed both striking repeatability and profound stochasticity, depending on the level of analysis^5^. Convergence—where distinct lineages independently evolve similar traits under analogous selective pressures—appears more widespread than previously appreciated^6^. Natural selection frequently yields comparable fitness gains even as populations retain the imprint of distinct genetic histories. However, laboratory systems often simplify ecological and genetic complexities, limiting their relevance to natural adaptation. Field-based studies thus serve as essential complements, capturing evolutionary dynamics in ecologically realistic contexts^7,8^. Comparative population genomics has further revealed both convergent and divergent responses across diverse taxa, including animals^9–14^, plants^15–18^, and pathogens^19^. Yet, many studies have been limited by small sample sizes or narrow geographic ranges. More critically, prior work has primarily focused on either long-term (species level)^14,20,21^ or short-term (population level)^12,13,15,16,19^ evolutionary timescales, thereby lacking a comprehensive perspective on the continuous and dynamic nature of adaptive evolutionary trajectories across multiple temporal scales. Moreover, convergence is rarely perfect: even under similar selective regimes, adaptive trajectories can diverge at the molecular level, shaped by differences in genetic architecture, demography, and evolutionary history. Therefore, conducting population genomic studies on closely related, globally distributed species with similar evolutionary histories allows for a comparative investigation of their environmental adaptation trajectories across both ancient and recent evolutionary timescales.

The *Drosophila* genus, known for its ecological breadth, provides a powerful system for studying the mechanisms of adaptation in nature^22–24^. Among *Drosophila*, *D. melanogaster* exhibits pronounced seasonal adaptation, repeatedly evolving similar phenotypes in response to predictable environmental fluctuations within and across years^7,8^. Moreover, *D. melanogaster* and its sibling species *D. simulans* present a particularly compelling system for studying evolutionary convergence. Diverging less than 5 million years ago, these sibling species share a common African origin and have independently colonized the diverse global environments^25–28^. Despite their ecological similarity and parallel history of range expansion, they differ in their genomic architecture, geographic distribution, population genetic features^29^ (e.g., genetic diversity, effective population size [*N*_e_], etc.), and key adaptive traits such as alcohol tolerance^30^ and thermal tolerance^29,31^. Both species have encountered comparable selective pressures—such as exposure to insecticides and climate variability—during their worldwide dispersal, providing a natural experiment for investigating the repeatability and contingency of adaptation under shared environmental challenges.

Here, we present a large-scale comparative analysis of globally sampled *D. melanogaster* and *D. simulans* populations. We assembled a comprehensive genomic dataset comprising 1,335 *D. melanogaster* genomes and 681 *D. simulans* genomes, including hundreds of newly sequenced strains from China. By tracing signatures of adaptation across continents, we identified both convergent and species-specific evolutionary responses—most notably to insecticide exposure. These findings provide a natural evolutionary replay experiment, offering insights into how and why evolution follows repeatable and predictable paths.

## Results

### High-quality chromosome-level genome assembly and annotation of *D. simulans*

Previous *D. simulans* assemblies—based on Sanger^32,33^, Illumina^34,35^, or hybrid PacBio– Illumina^36^ sequencing—have facilitated genetic studies but remain fragmented (e.g., the current reference dsim_r2.02 [Dsim2] has a contig N50 of 169 kb^37^). To improve contiguity and accuracy, we assembled a chromosome-level genome for the *w*^501^ strain using PacBio HiFi (∼120.18× coverage), ultralong ONT (∼ 322.34× coverage), and Hi-C data (∼187.44× coverage) (Figures 1A, S1A, and Table S1).

**Figure 1.**
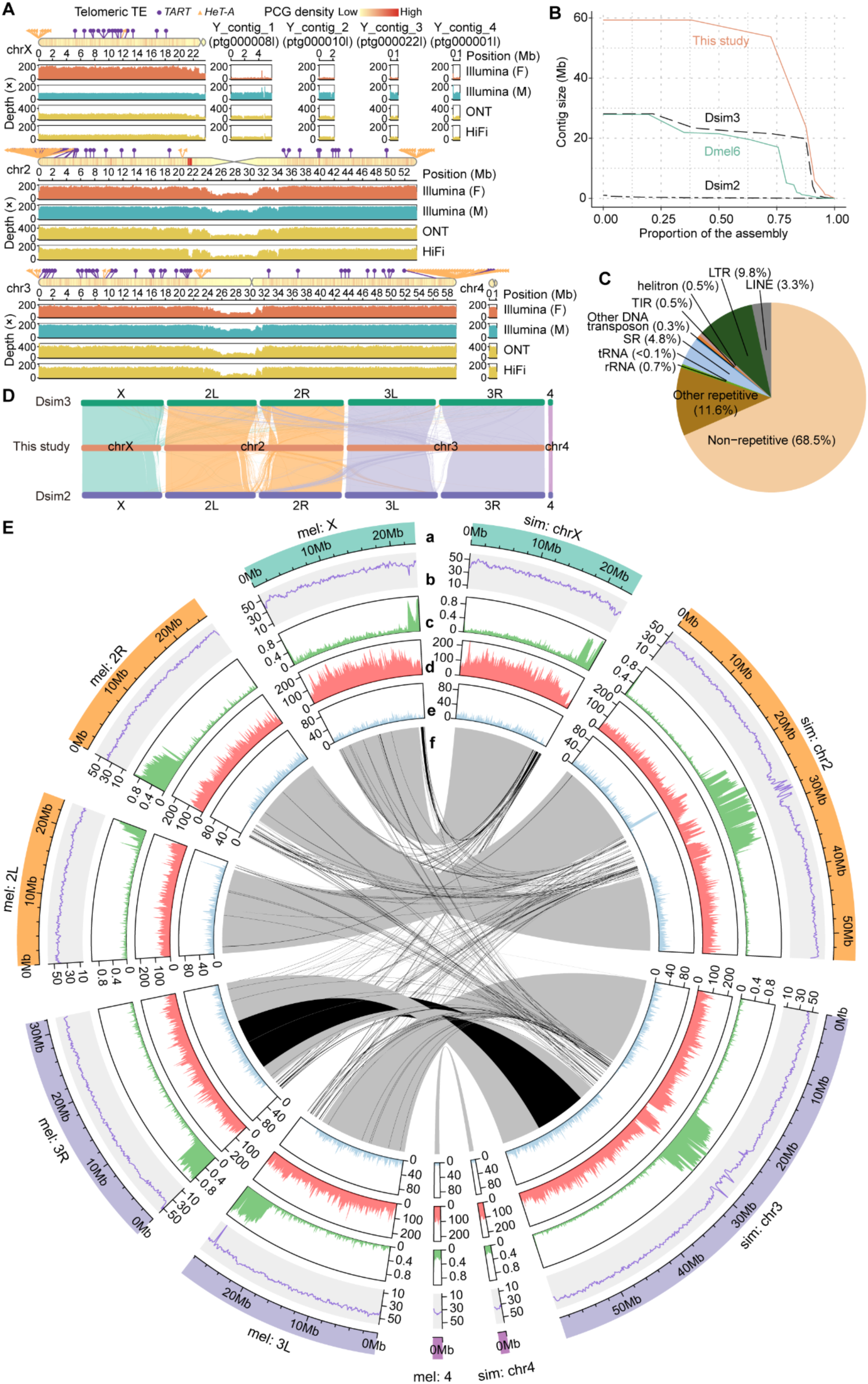
Genomic features and quality evaluation of the newly assembled *D. simulans* genome, and comparative genomic analysis with *D. melanogaster*. (A) Genomic features and sequencing depth of coverage of the *D. simulans* genome. Shown are chrX, chr2, chr3, chr4, and Y-linked contigs >1 Mb. In each panel, the first row depicts the chromosome schematic with telomeric transposable elements (*HeT-A*, yellow triangles; *TART*, purple circles). Gene density is color-coded from low (yellow) to high (red) in 100 kb nonoverlapping windows. The second row shows genomic coordinates, followed by sequencing depth of Illumina PCR-free libraries (virgin female [F] and male [M]), Oxford Nanopore Technology (ONT), and PacBio HiFi data. High mapping rates across PacBio HiFi (98.42%), ONT (95.64%), and Illumina reads (99.29% for males and 99.19% for females), together with genome continuity inspector (GCI) scores (chrX: 99.9999, chr2: 99.9585, chr3: 100.0, chr4: 99.9981), confirm chromosomal integrity and completeness. Four major chromosomes (chrX: 23.74 Mb; chr2: 53.73 Mb; chr3: 59.32 Mb; chr4: 1.16 Mb) and four Y-linked contigs (Y_Contig_1–4) are shown. (B) Nx curves showing the assembly contiguity of the genomes. The final assembly achieved a contig N50 of 53.75 Mb. The newly assembled *D. simulans* genome assembly (orange) was compared to two previous genome versions of *D. simulans* (Dsim3 and Dsim2, black), as well as the *D. melanogaster* r6.54 genome (Dmel6, green). The *y*-axis represents the size of each contig in descending order, whereas the *x*-axis shows the cumulative proportion of the genome assembly that is covered. (C) Proportion of repetitive sequences in the newly assembled genome of *D. simulans*. The genome of *D. simulans* is composed of 31.5% repetitive and 68.5% nonrepetitive sequences. (D) Synteny analysis showing a one-to-one correspondence between the four chromosomes of different *D. simulans* genome assemblies, including Dsim3 from NCBI (top), the genome assembly from the present study (middle), and Dsim2 from FlyBase (bottom). Each line represents an orthologous block between the compared genomes, color-coded according to their chromosome of origin. (E) Comparison of genomic features between *D. melanogaster* (left) and *D. simulans* (right). The rings from outside to inside indicate the genomic coordination (a), GC content percentage of 100-kb nonoverlapping windows (b), percentage of repeat elements per 100-kb nonoverlapping window (c), number of single nucleotide polymorphisms (SNPs) (d) and genes (e) per 100-kb nonoverlapping window, and syntenic blocks shared between *D. simulans* and *D. melanogaster* (f). The color represents inverted (black) and noninverted regions (grey).

The final 156.08 Mb assembly achieved a contig N50 of 53.75 Mb, comprising four major chromosomes, a complete mitochondrial genome, and 23 unplaced contigs, including 13 confidently assigned to the Y chromosome (Figures 1B, S1B and Table S2). Quality assessment confirmed high accuracy, completeness (BUSCO: 99.06%), and uniform coverage across platforms (Figure S1C, and Tables S3–S4). Repetitive sequences constituted 31.54% (49.22 Mb) of the genome, with LTRs representing the largest repeat class (Figure 1C and Table S5). We annotated 14,927 protein-coding genes along with diverse noncoding RNAs, surpassing previous *D. simulans* assemblies and even *D. melanogaster* (Tables S6–S8). Synteny analysis verified chromosome integrity, corrected previous misassemblies, and validated known structural variants such as *In(3R)84F1;93F6–7* in *D. melanogaster* (Figures 1D–E and S1D).

Together, this chromosome-level assembly provides a highly accurate and complete reference, establishing a valuable resource for population and evolutionary genomics of *D. simulans*.

### Contrasting genomic diversity landscapes in *D. simulans* and *D. melanogaster*

To investigate global patterns of diversity and divergence, we analyzed whole-genome sequences from 1,335 *D. melanogaster* and 681 *D. simulans* strains (Tables S9–S10). In *D. melanogaster*, 108 new iso-female strains from China were sequenced and integrated with 292 previously published genomes^38^, yielding 400 genomes from China (Figure 2A). Combined with 935 published genomes^28,39–50^, this dataset comprised 1,335 genomes across 22 populations (Figure 2B). For *D. simulans*, we sequenced 419 new strains from China and combined them with 262 published genomes, generating 681 genomes from 11 populations spanning Africa^51–55^, Europe^56^, North America^57^, and Asia^58^ (Figure 2A–B). The 537 newly sequenced strains had mean coverages of 25.1× (*D. melanogaster*) and 21.4× (*D. simulans*) (Table S9). This dataset enabled comparative assessment of convergent and divergent evolutionary trajectories in these sibling species.

**Figure 2.**
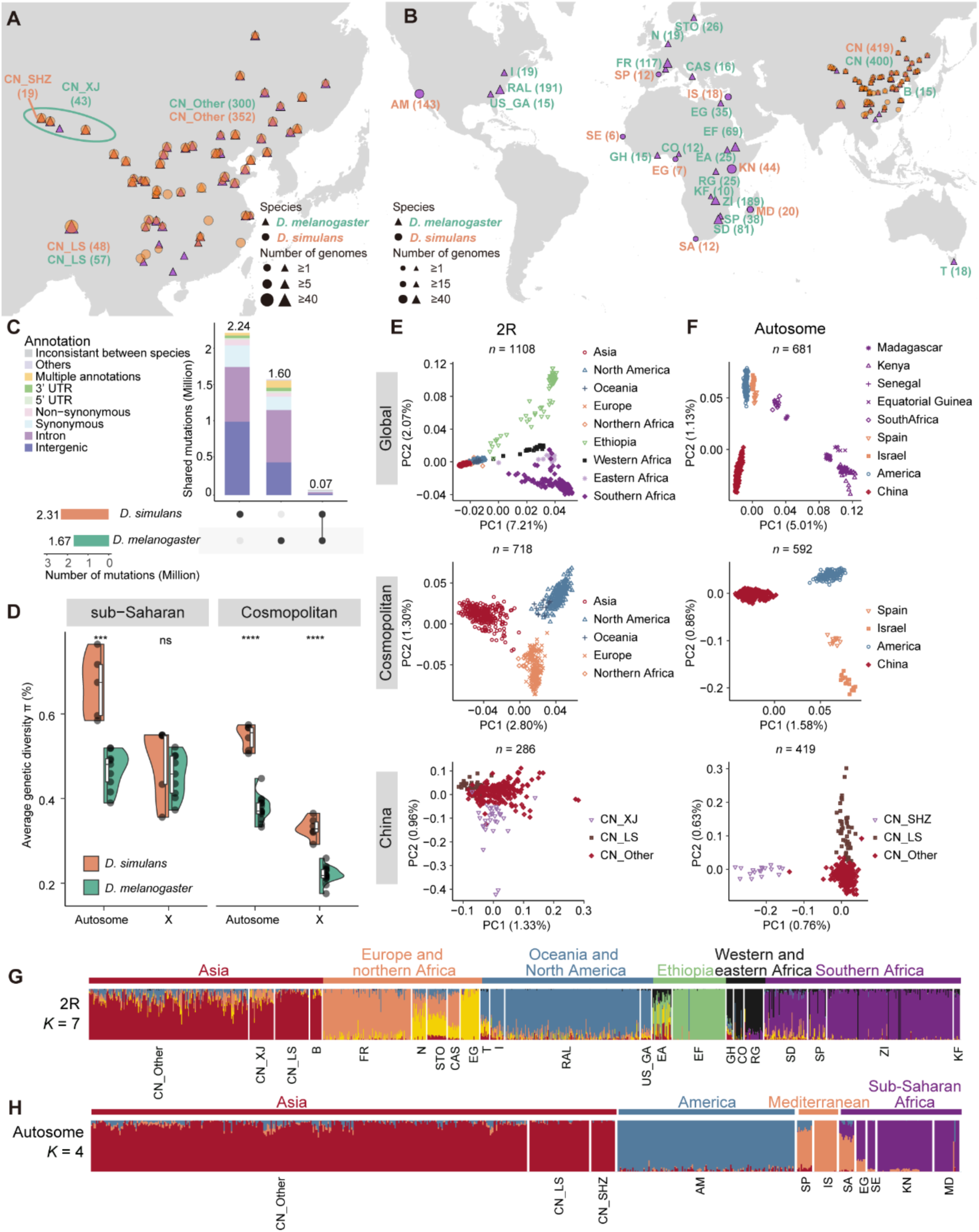
Sample locations, genetic diversity and population structure of global *D. simulans* and *D. melanogaster* populations. (A) Geographic locations of 419 *D. simulans* (circles) and 400 *D. melanogaster* (triangles) strains collected across China (CN). *D. melanogaster* strains were sampled from Lhasa, Tibet (CN_LS, *n* = 57); Xinjiang (CN_XJ, *n* = 43; including Hami, Turpan, Shihezi, and Urumqi); and other regions in China (CN_Other, *n* = 300). *D. simulans* strains were sampled from Lhasa, Tibet (CN_LS, *n* = 48); Xinjiang (Shihezi [CN_SHZ], *n* = 19; Hami and Urumqi, *n* = 21); and other regions (CN_Other, *n* = 331). For *D. melanogaster*, orange symbols represent the 108 newly sequenced strains, and purple symbols indicate previously published genomes. All the *D. simulans* strains were newly sequenced in this study. (B) Sample locations of 681 *D. simulans* genomes and 1,335 *D. melanogaster* genomes involved in this study. More details are given in Tables S9 and S10. (C) Overlap of common SNPs (MAF > 0.05) identified in *D. simulans* and *D. melanogaster*. Stacked bar graphs showing the number of SNPs of different annotation types. The results for all the polymorphic sites are shown in Figure S2A. (D) Split violin plots showing the genetic diversity of *D. simulans* (orange) and *D. melanogaster* (green). The average π was calculated on the basis of all strains of each population of *D. simulans* and *D. melanogaster*. The top and bottom of the boxes within the split violin depict the 75^th^ and 25^th^ percentiles of the distribution. The horizontal lines within the boxes signify the median values. Axis endpoints are labeled by the minimum and maximum values. The statistical significance according to the Wilcoxon rank sum test is shown for each comparison. ns: *P* > 0.05, ***: *P* < 0.001, ****: *P* < 0.0001. (**E** and **F**) Principal component analysis (PCA) results of *D. melanogaster* (**E**) and *D. simulans* (**F**). The first two principal components (PCs) were plotted for global strains, cosmopolitan strains, and strains in China, respectively. The proportion of variance explained by each corresponding principal component is indicated in parentheses, with the number of genomes (*n*) labeled at the top of the graph. The results are based on neutral sites from inversion-free *D. melanogaster* strains on 2R and *D. simulans* strains on autosomes (combining chr2, chr3, and chr4). (**G** and **H**) Admixture proportions inferred for each strain in 22 populations of *D. melanogaster* (**G**) and 11 populations of *D. simulans* (**H**), with CN subpopulations shown separately. Strains are sorted by their geographic origin and represented by color-coded vertical columns that reflect the compositions of their ancestry. The results, which are based on neutral SNPs in inversion-free 2R of *D. melanogaster* and autosomes of *D. simulans* are displayed. The optimal number of ancestries (*K*) was determined when the minimum cross validation (CV) error median was reached (Table S13).

Although large inversions are common in *D. melanogaster*^38,59^ (Table S11), none have been reported in *D. simulans*. Using a unified variant-calling pipeline, we identified ∼13.82 million single nucleotide polymorphisms (SNPs) in *D. melanogaster* and ∼10.93 million in *D. simulans* (Figure 1E, and Table S12). Among them, 12.3% of *D. melanogaster* SNPs and 21.4% of *D. simulans* SNPs were common variants (minor allele frequency [MAF] > 0.05), indicating more frequent common alleles in *D. simulans* (Figure 2C).

Alignment of the new *D. simulans* reference to *D. melanogaster* improved coverage of orthologous sites relative to earlier assemblies (Figure S1E–F). Among *D. melanogaster* SNPs, 97.1% mapped to the new *D. simulans* genome, with 2.16 million sites polymorphic in both species (19.7% of *D. simulans* and 15.6% of *D. melanogaster* SNPs; Figure S2A). Most shared polymorphisms were rare (MAF < 0.05), with only 72,126 (3.33%) common in both species, and the majority located in synonymous or noncoding regions (Figure 2C).

Across populations, autosomal diversity (π_A_) was significantly higher in *D. simulans* than in *D. melanogaster* in both sub-Saharan African (median: 0.674% vs. 0.459%, *P* < 0.001, Wilcoxon rank sum test) and cosmopolitan populations (median: 0.544% vs. 0.370%, *P* < 0.001, Wilcoxon rank sum test) (Figures 2D and S2B), consistent with the larger *N*_e_ in *D. simulans*. For X chromosome, *D. simulans* showed higher, but non-significant, nucleotide diversity (π_X_) than *D. melanogaster* in sub-Saharan Africa (median: 0.548% vs. 0.458%, *P* > 0.05), and significantly higher π_X_ values in cosmopolitan populations (median: 0.322% vs. 0.218%, *P* < 0.0001, Wilcoxon rank sum test). These results highlight substantial contrasts in genome-wide variation between the two species, reflecting differences in mutation bias, demography, and chromosomal context.

### Population structure and chromosomal differentiation in *D. simulans* and *D. melanogaster*

To account for demographic effects on genome-wide selection scans^60,61^, we analyzed population structure in both species using SNPs from putatively neutral regions (short introns and fourfold degenerate sites^62^). Strains with common large inversions in *D. melanogaster* were excluded from principal component analysis (Table S11), while all *D. simulans* strains were included. Both species showed strong differentiation between sub-Saharan and cosmopolitan populations (Figures 2E–F and S2C–D). Within Africa, *D. melanogaster* formed three clusters (southern, western/eastern, Ethiopian), mirrored by *D. simulans* (southern Africa, western/eastern Africa, Madagascar). Cosmopolitan populations grouped into three regions in each species: North America/Oceania, Europe/Northern Africa, and Asia for *D. melanogaster*; and North America, Mediterranean (Israel [IS] and Spain [SP]), and Asia for *D. simulans*. Strains from China (CN) formed distinct clusters within cosmopolitan groups in both species.

ADMIXTURE^63^ analyses supported these patterns (Table S13). In *D. melanogaster*, six ancestries were inferred: three sub-Saharan groups (southern Africa, western/eastern Africa, Ethiopia) and three cosmopolitan groups (Europe/Northern Africa, America/Oceania, Asia) (Figures 2G and S2E). In *D. simulans*, the best-fit model clearly separated sub-Saharan Africa from three cosmopolitan groups: North America, Mediterranean, and Asia (Figures 2H and S2F).

Fine-scale analyses revealed strong differentiation within strains from China. In *D. melanogaster*, PCA distinguished Tibet (CN_LS), Xinjiang (CN_XJ), and other strains from China (CN_Other) (Figures 2E and S2C). In *D. simulans*, strains from Shihezi, Xinjiang (CN_SHZ, *n* = 19) formed a distinct cluster along PC1, while those from Hami and Urumqi grouped with CN_Other. Differentiation between Tibet (CN_LS) and CN_Other was evident along PC2 (Figures 2F and S2D). ADMIXTURE (*K* = 6) supported distinct ancestries for CN_SHZ and CN_LS, with chromosome-specific patterns (Figure S2G). We propose that CN_SHZ strains represent an endemic Xinjiang lineage, whereas Hami and Urumqi strains likely reflect recent immigration; however, local differentiation within CN_SHZ cannot be excluded.

Together, these results reveal broadly shared population structure in the two species, with strong sub-Saharan versus cosmopolitan divergence and fine-scale chromosomal differentiation within China, reflecting demographic history and local adaptation.

### Population relationships and gene flow in *D. melanogaster* and *D. simulans*

TreeMix^64^ analyses of nine *D. simulans* populations (>10 strains each) identified optimal migration models of *m* = 2 for autosomes and *m* = 1 for the X chromosome. Rooted with Madagascar (MD)^26,27^, trees showed consistent topologies across chromosomes: Kenya (KN) and MD occupied basal positions, South African (SA) was near the root, and populations from China (CN_SHZ, CN_LS, CN_Other) clustered at the tips, closely related to American (AM), followed by Mediterranean populations (IS and SP) (Figures 3A and S3A).

**Figure 3.**
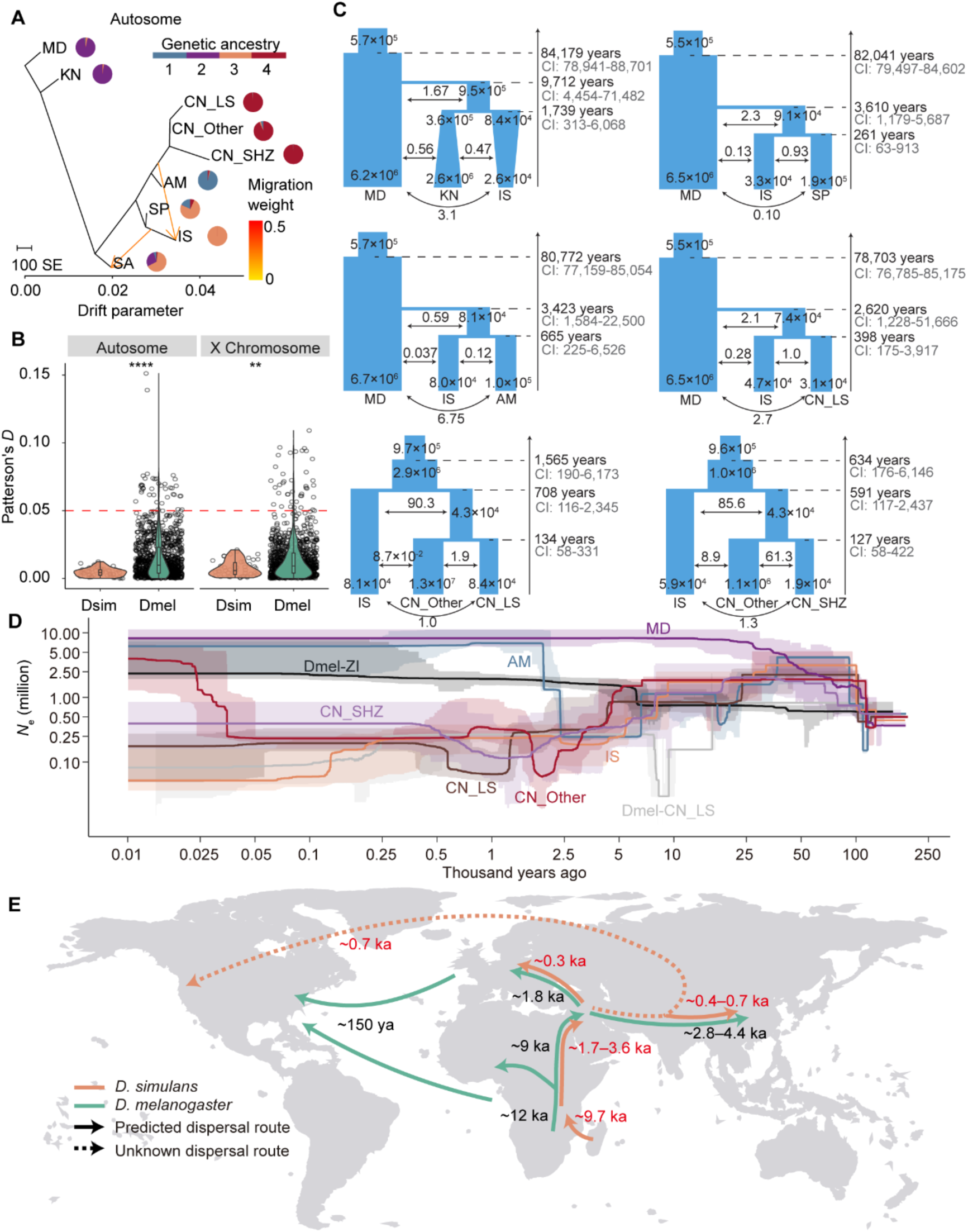
Gene flow and demographic history of *D. simulans* and *D. melanogaster*. (A) The maximum likelihood tree inferred using TreeMix based on nine populations of *D. simulans* with sequenced genome samples larger than 10. The population names were followed by the genetic ancestry compositions inferred by ADMIXTURE. The analysis was based on neutral sites of autosomes for *D. simulans*. The optimal number of migration edges (*m*) was determined via the OptM package. Two migration edges were inferred in *D. simulans*: 1) from the ancestor of CN to IS and 2) from the ancestor of IS to SA. Populations include Madagascar (MD), Kenya (KN), Israel (IS), Spain (SP), American (AM), South African (SA), and China (CN). (B) Comparison of Patterson’s *D* values of *D. simulans* and *D. melanogaster* on autosomes and X chromosomes. The red dashed line represents the threshold of *D* > 0.05 used. Statistical significance was determined using the Wilcoxon rank sum exact test for each comparison, **: *P* < 0.01, ****: *P* < 0.0001. (C) Best-fitting evolutionary models involving the *D. simulans* populations inferred by moments based on neutral SNPs of autosomes. Parameters such as the effective population size (*N*_e_), split time, and migration strength (2*N_m_*) are shown. (D) Demographic history of six representative populations (MD, AM, IS, CN_SHZ, CN_LS and CN_Other) of *D. simulans* using the unfolded site frequency spectrum (SFS) of neutral sites on autosomes, as analyzed by Stairway Plot 2. Historical effective population size (*N*_e_) changes from 0.01 to 200 ka are depicted. The solid lines represent the medians of the inferred population sizes, whereas the light ribbons indicate the 95% confidence intervals. A mutation rate of 4.51 × 10^-9^ per site per generation and a generation time of 15 per year were used. Two additional curves analyzed in a previous study^38^ represent ZI and CN_LS of *D. melanogaster*, which are shown in black and gray, respectively. (E) Proposed range expansion and migration routes of *D. simulans* (green) and *D. melanogaster* (orange). The predicted expansion routes are represented by solid arrows, and expansion routes that cannot be determined are indicated as dashed lines. Time estimates were estimated in this study (in red) or previous studies (in black). ya: years ago; ka: kilo annum.

While *D. melanogaster* exhibits extensive genetic admixture^38,41,65^ (Figures 3B and S3B–D), gene flow in *D. simulans* appeared limited. Three possible migration events were inferred in *D. simulans* by Treemix: autosomal gene flow from CN to IS and IS to SA, and X-linked gene flow from SA to CN. The admixture graph analyses suggested two additional admixtures: (1) 87.5% Mediterranean ancestry and 12.5% African ancestry in the AM and CN populations, and (2) a minor North American/Asian contribution (6.1%) to MD (Figure S3E). However, Patterson’s *D* statistics^66^ across 46 population trios detected no significant gene flow (|*D*| > 0.05; FDR < 0.05) in *D. simulans* (Table S14). Together, these results point to weak or inconsistent signals across methods, reinforcing that gene flow is far more restricted in *D. simulans* than in *D. melanogaster*.

*D. simulans* exhibited consistently higher *D*_XY_ values than *D. melanogaster* across both cosmopolitan and sub-Saharan African populations (Figure S3F), likely reflecting greater ancestral diversity and demographic history. Unlike *D. melanogaster*, which showed isolation by distance^38^, *D. simulans* did not, suggesting differences in demographic history or recent long-distance migrations facilitated by human activities (Figure S3G–H and Table S15). These findings also support the idea that *D. melanogaster* is more closely associated with humans, whereas *D. simulans* is not a strictly human commensal^26^.

### A more recent and rapid global expansion of *D. simulans* compared to *D. melanogaster*

Despite its presumed origin in Madagascar^26,27^, the global demographic history of *D. simulans* remains incompletely defined. Using neutral SNPs, we inferred the population divergent times and effective population size (*N*_e_) trajectories of *D. simulans* (Figure 3C–D and Table S16). The MD population diverged from the KN–IS ancestor ∼9.7 thousand years ago (ka) (95% CI: 4.5–71.5), indicating early African expansion. KN and IS then split ∼1.7 ka (95% CI: 0.3–6.1), marking the out-of-Africa migration. Cosmopolitan populations colonized more recently: SP ∼261 years ago (95% CI: 63–913), AM ∼665 years ago (95% CI: 225–6,526), and CN ∼398 years ago (95% CI: 175–3,917). Within China, CN_SHZ and CN_LS diverged from CN_Other ∼127 (95% CI: 58–422) and ∼134 years ago (95% CI: 58–331), respectively.

Compared with *D. melanogaster*, which dispersed from Africa ∼9 ka^38^, *D. simulans* expanded later (∼1.7–3.6 ka) (Figure 3E). Its arrival in East Asia (∼0.3 ka) and Europe (∼0.4–0.7 ka) lagged behind *D. melanogaster* (∼1.8 and 2.8–4.4 ka, respectively). In contrast, *D. simulans* colonized America earlier (∼700 years ago vs. 150 years ago for *D. melanogaster*^67^), though no historical records support this estimate. Across time, *D. simulans* consistently exhibited higher *N*_e_, about fivefold greater than *D. melanogaster*. For example, ∼400 years ago, *N*_e_ in MD reached ∼8 × 10⁶, compared with ∼1.2 × 10⁶ in *D. melanogaster* from Zambia (ZI)^38^. Overall, these results highlight the more recent but rapid global expansion of *D. simulans*, its higher ancestral diversity, and distinct demographic history, in contrast to the earlier, more bottlenecked spread of *D. melanogaster*.

### Convergent adaptation in the long-term divergence of *D. melanogaster* and *D. simulans*

To investigate long-term adaptive protein evolution between *D. melanogaster* and *D. simulans*, we applied the McDonald–Kreitman test (MKT)^68^, using *D. yakuba* as an outgroup (Figure 4A). Fixed nonsynonymous and synonymous substitutions were compared to polymorphic variants, and *α*, the proportion of fixed amino acid changes driven by positive selection, was estimated. Genes with at least six mutations per category were retained for analysis^14,69^.

**Figure 4.**
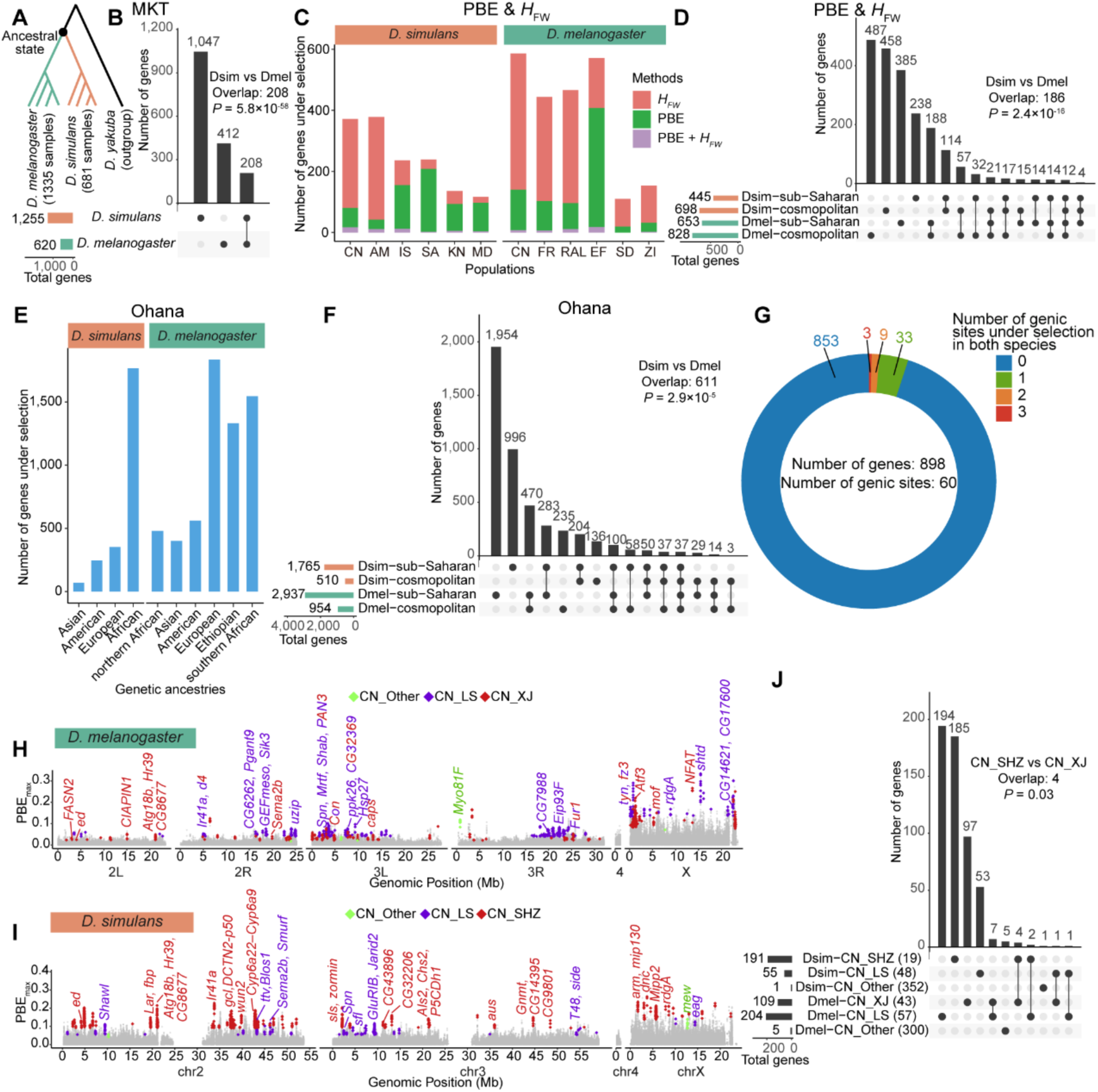
Convergent adaptation between *D. simulans* and *D. melanogaster*. (A) Schematic drawing of the phylogenetic tree for McDonald–Kreitman test (MKT). The outgroup *D. yakuba* was used to estimate the ancestral state at the node of the most recent common ancestor (MRCA) of *D. simulans* and *D. melanogaster*. (B) Overlap of positively selected genes detected by MKT in *D. simulans* and *D. melanogaster*. The number of positively selected genes (*P* < 0.05) in the *D. melanogaster* lineage (412), the *D. simulans* lineage (1,047), and both (208), are shown. The statistical significance was conducted using the hypergeometric test. (C) The number of genes showing a selection signal at the population level according to Fay and Wu’s *H* (*H*_FW_) and population branch excess (PBE) in *D. simulans* and *D. melanogaster* populations. *D. simulans* populations include China (CN), American (AM), Israel (IS), South African (SA), Kenya (KN), and Madagascar (MD); and *D. melanogaster* populations include China (CN), France (FR), America (RAL), Ethiopia (EF), South Africa (SD), and Zambia (ZI). (D) Overlap of positively selected genes detected by *H*_FW_ and PBE in *D. simulans* and *D. melanogaster*. Populations were grouped as sub-Saharan or cosmopolitan based on geographic origin, with three populations per group. Statistical significance of the overlap was assessed using a hypergeometric test. (E) The number of genes showing selection signals at the level of genetic ancestry by Ohana in *D. simulans* and *D. melanogaster* populations. (F) Overlap of positively selected genes detected by Ohana in *D. simulans* and *D. melanogaster*. The genetic ancestries of each species were grouped according to their geographic locations. Statistical significance of the overlap was assessed using a hypergeometric test. (G) Classification of 898 adaptively convergent genes based on the number of shared SNPs under selection in both species. A total of 87 shared SNPs (60 genic) were under convergent selection in both species, including 3 genes with 3 SNPs, 9 genes with 2 SNPs, and 33 genes with a single SNP. (**H** and **I**) Manhattan plots showing branch-specific positive selection detected by PBE in three subpopulations from China in *D. melanogaster* (**H**) and *D. simulans* (**I**). For each subpopulation, the max PBE values of two trio tests (see Figure S9C–D) are shown for each 4-kb sliding window with a step size of 1 kb. The candidate windows are colored according to population, and genes potentially related to environmental adaptation and mentioned in the main text are annotated. Gene names with mosaic colors represent genes that are selected from multiple populations. (**J**) Overlap of positively selected genes detected by PBE in three subpopulations from China of *D. simulans* and *D. melanogaster*. The number of genomes used in the analysis for each subpopulation is indicated in parentheses. In *D. simulans*, strains from Shihezi, Xinjiang (CN_SHZ) likely represent an endemic Xinjiang lineage, whereas strains from Hami and Urumqi were grouped with CN_Other, possibly reflecting recent migration into the region. The statistical significance based on the hypergeometric test is shown.

In the lineage leading to extant *D. melanogaster*, 5,790 genes were analyzed; 620 (10.7%) showed nominal evidence of positive selection (*P* < 0.05), with an overall *α* = 0.77 (Figure 4B and Table S17). After FDR correction, 80 genes remained significant (*α* = 0.81). The 620 selected genes were enriched in dosage compensation and meiotic cell cycle pathways (Figure S4A and Table S18). Notable examples include genes involved in TOR signaling (*raptor* and *MAPK-Ak2*), immunity (*spirit*, *TotB*, and *RluA-2*), stimulus detection (*Pepck1*, *Gr63a*, and *Ir7c*), and courtship behavior (*ppk29*, *tilB*, and *Gr10b*). Interestingly, selected genes were significantly enriched on the X chromosome (166 X-linked vs. 454 autosomal; *P* < 0.001, Fisher’s exact test), supporting the faster-X hypothesis^70–73^.

In the *D. simulans* lineage, 5,614 genes were analyzed, with 1,255 (22.4%) showing nominal signatures of positive selection (*P* < 0.05), and 527 genes significant after FDR correction (overall *α* = 0.85; Figure 4B). These 1,255 genes were enriched in reproductive and immune pathways (Figure S4B), such as *mmm* and *lobo* (sperm function), *Mrp4* and *Daxx* (lifespan regulation), *Tep1* and *Tak1* (immune response), and *Mtr4* and *Rrp6* (virus defense). Unlike in *D. melanogaster*, no enrichment was found on the X chromosome (165 X-linked vs. 1,090 autosomal; *P* > 0.05). Notably, genes in DNA repair pathway showed stronger positive selection signals in *D. simulans* (e.g., *Nup160*, *PolZ1*, *Marcal1*, *Rad9*, *Irbp*, and *Rad60*) compared to *D. melanogaster*, possibly explaining the higher chromosomal inversion occurrences in *D. melanogaster*.

Strikingly, 208 positively selected genes were shared between species, more than expected by chance (*P* = 5.79 × 10^−58^, hypergeometric test), enriched in reproductive and genome defense pathways (Figure S4C). Shared genes include *Pkd2* (sperm storage), *Dhc16F* and *mip120* (sperm motility), *Dnaaf3* (sperm assembly), and piRNA pathway genes (*aub*, *fs(1)Yb*, *qin*, and *tud*), highlighting convergent adaptation in male reproduction and suppression of transposable elements.

### Ancient signals of pathway-level convergent adaptation in populations of the two sibling species

To investigate adaptive signals within populations of *D. melanogaster* and *D. simulans*, we applied Fay and Wu’s *H* (*H*_FW_) test^74^, which detects an excess of high-frequency derived alleles—indicative of positive selection maintained across broad geographic and demographic populations. We analyzed six geographically matched populations of each species (*D. melanogaster*: CN, FR, RAL, EF, SD, ZI; *D. simulans*: CN, AM, IS, SA, KN, MD), scanning 4-kb windows (1-kb steps). Outlier regions, defined as the lowest 1% of the species-specific *H*_FW_ distribution (*H*_FW_ < −0.53 for *D. melanogaster*, *H*_FW_ < −0.28 for *D. simulans*; Figure S4D– F), identified regions shaped by repeated or widespread selection at the population level.

We identified 544 genes with signatures of positive selection in at least one *D. melanogaster* population and 478 in *D. simulans* (Figure S4G–H and Table S19). In *D. simulans*, 98% of selected genes occurred in cosmopolitan populations (309 in CN, 347 in AM, 93 in IS), with only nine unique to sub-Saharan Africa. Seventy genes showed signatures in both species (Figure S4I), enriched for pathways including insecticide metabolism, alcohol biosynthesis, and DDT response (Figure S4J).

Cross-species comparison revealed convergent adaptation at the pathway level between the two species, notably in “RNA processing” and “insecticide catabolic process” pathways (Figure S4K–L). While “RNA processing” was enriched within each species, only two genes (*Adar* and *SmD3*) overlapped between species. *Adar*, a key RNA-editing enzyme^75^, was selected in the CN populations of both species and in additional *D. simulans* (AM, KN, and MD) and *D. melanogaster* (FR) populations, suggesting its role in environmental adaptation.

These results indicate that convergent selection acts primarily at the pathway level— particularly in environmental stress and insecticide response—while adaptive responses at the gene level remain largely species-specific, reflecting lineage-specific constraints and historical contingency.

### Population-specific adaptation and recent convergence in the two species

To identify recent, population-specific adaptations, we applied population branch excess (PBE) analysis^76^ to six representative populations of *D. melanogaster* and *D. simulans* (Figure S5A– B). Unlike *H*_FW_, which typically detects older selection signals shared across populations, PBE identifies recent selective sweeps in focal populations. Accordingly, we observed limited overlap between genes identified by PBE and *H*_FW_ within each population (Figure 4C). In *D. melanogaster*, PBE identified 1,045 candidate genes (140 in CN, 102 in FR, 96 in RAL, 408 in EF, 19 in SD, and 32 in ZI; Figure S5C–D), consistent with previous findings^38^. In *D. simulans*, 620 candidate genes were detected (97 in MD, 93 in KN, 208 in SA, 155 in IS, 42 in AM, and 80 in CN; Figure S5E–F).

Despite the population-specific nature of PBE, 94 genes were detected in both species, suggesting convergent responses to similar selective pressures (Figure S5G). These genes were enriched in pathways related to the chemical stimulus response, developmental growth regulation, autophagy, and synaptic transmission (Figure S5H). Examples include *Ace* (insecticide resistance), under selection in *D. simulans* IS and *D. melanogaster* CN; *mTor*, a metabolic regulator, selected in *D. simulans* MD and *D. melanogaster* EF; and RNAi genes *AGO2* and *AGO3*, under selection in *D. simulans* KN and *D. melanogaster* FR/ZI, respectively (Table S19). These results indicate that as *D. simulans* and *D. melanogaster* colonized diverse environments, they experienced similar selective pressures that drove convergent adaptation at both the gene and pathway levels—although through largely independent, population-specific trajectories (Figure 4D).

### Ongoing convergent selection in populations of *D. melanogaster* and *D. simulans*

While both *H*_FW_ and PBE can be confounded by demography, we applied the Ohana framework^77,78^, which accounts for population structure to detection of positive selection at the ancestry level (Figure S6A–B). Ohana identified 3,270 candidate genes under positive selection in *D. melanogaster* (1,545 in southern African, 1,331 in Ethiopian, 1,834 in northern African, 562 in European, 479 in Asian, 400 in North American ancestries; Figures 4E and S6C) and 1,947 in *D. simulans* (68 in Asian, 245 in American, 352 in Mediterranean, 1,765 in sub-Saharan African; Figure S6D). Of these, 611 genes were shared between species, significantly more than expected (*P* = 2.94 × 10⁻⁵, hypergeometric test; Figure 4F), enriched primarily in sub-Saharan ancestries and developmental/morphogenetic pathways (Figure S6E).

Cross-method comparison revealed limited overlap among genes identified by Ohana, *H*_FW_, and PBE. In *D. melanogaster*, 1.42–20% of Ohana hits overlapped with either method; in *D. simulans*, the overlap ranged from 8.32–25.57% (Figure S7A–B). Integrating all three approaches yielded 898 genes under convergent selection between species (Figure S7C), encompassing 10,118 SNPs putatively selected in *D. simulans* and 14,265 in *D. melanogaster*. However, only 87 SNPs (60 in the genic regions) at orthologous positions were shared between species (Figure 4G), including just three nonsynonymous mutations: *pcm* (A1515T in *D. melanogaster*), involved in spermatogenesis and mRNA decay^79,80^; *CG6933* (T218P), linked to chitin-binding and melanism^81,82^; and *Cyp6g1* (T316S), associated with DDT resistance^83,84^ (Figure S8).

Collectively, our integrative analyses spanning divergence-based (MKT), population-level (*H*_FW_, PBE), and ancestry-aware (Ohana) approaches reveal a layered landscape of adaptive evolution in *D. melanogaster* and *D. simulans*. While we observed substantial convergence at the pathway and gene levels—particularly in functions related to reproduction, immunity, and the environmental response—convergence at the individual site level was rare. These findings underscore how shared selective pressures can drive parallel evolutionary outcomes through distinct molecular routes, shaped by species-specific demography, genetic architecture, and historical contingency.

### Local and convergent selection in the CN subpopulations of both species

By integrating selective signals from *H*_FW_, PBE and Ohana, we identified 39 genes showing convergent adaptation in the CN populations of both species (Figure S9A), enriched particularly in insecticide response pathways (e.g., *Cyp6v1*, *Cyp6g1*, *Cyp6g2*, and *Mdr65*; Figure S9B). To further characterize local adaptation, we applied PBE analysis to geographic subpopulations: CN_LS, CN_XJ, and CN_Other for *D. melanogaster*; and CN_LS, CN_SHZ, and CN_Other for *D. simulans*, using FR (*D. melanogaster*) and IS (*D. simulans*) populations as outgroups, respectively (Figure S9C–D).

In *D. melanogaster*, PBE detected 204 selected genes in CN_LS, 109 in CN_XJ, and five in CN_Other (Figure 4H). CN_LS candidates included circadian (*CG7988, Shab*), starvation resistance (*Hsp27*), mechanical sensing (*Ir41a, ppk26*), and hypoxia response (*Mrtf, Eip93F*) (Figure S9E). CN_XJ candidates involved DNA repair (*mof*), oxidative stress (*Atg18b*), nutrient sensing (*FASN2*), immune signaling (*Atf3*), and salt stress (*NFAT*).

In *D. simulans*, 191 genes were identified in CN_SHZ, 55 in CN_LS, and one (*mew*) in CN_Other (Figure 4I). CN_SHZ candidates included sugar metabolism (*fbp*), gamete generation (*Lar, gcl, wun2, DCTN2-p50*), and oxidative stress (*Sps1*) (Figure S9F). Strong signals were also observed at insecticide loci, including *Mdr50* and a *Cyp6a* gene cluster (*Cyp6a9, Cyp6a17, Cyp6a19, Cyp6a22, Cyp6a23*). CN_LS genes were enriched for synaptic vesicle recycling (*sfl, ttv, Blos1*) and potassium ion transport (*Shawl, GluRIB*) (Figure S9G).

Eight genes showed selection in CN subpopulations of both species, with five detected in parallel environments (Figure 4J). For example, *Spn* (synaptic regulation) was under selection in both Lhasa populations, while *ed* (cell differentiation), *CG8677* (histone exchange), *Atg18b* (oxidative stress), and *Hr39* (transcriptional regulation) were selected in Xinjiang populations of both species. This overlap was greater than expected by chance (*P* = 0.03, hypergeometric test; Figure 4J). These findings reveal both local adaptation and convergent evolution in CN strains of *D. melanogaster* and *D. simulans*, shaped by shared pressures such as hypoxia, temperature, and insecticide exposure across geographically structured populations.

### Parallel population cage experiments provide empirical evidence supporting cross-species genomic convergence

To empirically test whether different species utilize similar genetic mechanisms for convergent phenotypic outcomes, we performed cage-based selection experiments on oxidative stress resistance in *D. melanogaster* and *D. simulans*. For each species, ∼2,000 males (10 per strain from ∼200 strains collected across 20 locations from China; Table S20) were pooled and exposed to paraquat, a neurotoxic herbicide that induces oxidative stress^85^. From two biological replicates, we collected ∼200 resistant and ∼200 sensitive flies per replicate for whole-genome sequencing.

Genome-wide association using Cox proportional hazards modeling identified 1,584 SNPs in *D. melanogaster* and 632 SNPs in *D. simulans* (FDR < 5 × 10^-8^), mapping to 700 and 217 genes, respectively (Figure S10A–C). Differences in SNP and gene counts likely reflect detection thresholds rather than biological disparity. Enriched gene ontology categories differed between species (five terms in *D. melanogaster*, seven in *D. simulans*; Figure S10D–E). Among the 700 *D. melanogaster* candidates, 29 overlapped with previously identified oxidative stress genes^86,87^. Although no SNPs overlapped between species, 39 candidate genes were shared—significantly more than expected (*P* = 9.35 × 10^-11^, hypergeometric test)—indicating that convergent adaptation is more frequent at the gene level than at individual nucleotide sites (Figure S10C).

### Adaptive convergence in insecticide resistance in *D. melanogaster* and *D. simulans*

Multiple analyses consistently revealed enrichment of insecticide resistance pathways in both species, we compared positive selection signals in known resistance genes to investigate this further (Figure 5A–B). We identified 48 positively selected resistance genes in *D. melanogaster* populations (16 in cosmopolitan groups, 19 in sub-Saharan, 13 shared) and 36 in *D. simulans* (11 cosmopolitan, 14 sub-Saharan, 11 shared; Figure 5A). Overall, 20 of 119 resistance genes showed convergent selection between species. Convergence was especially pronounced in CN populations and ancestries, where 10.61% of resistance genes (7/66) were under selection in both species, compared to only 1.62% (21/1,300) in just one species (*P* = 0.0075, hypergeometric test). Notably, the *Cyp6g1–Cyp6t3* cluster showed recurrent selection across multiple ancestries in both species, underscoring its central role in convergent insecticide resistance.

**Figure 5.**
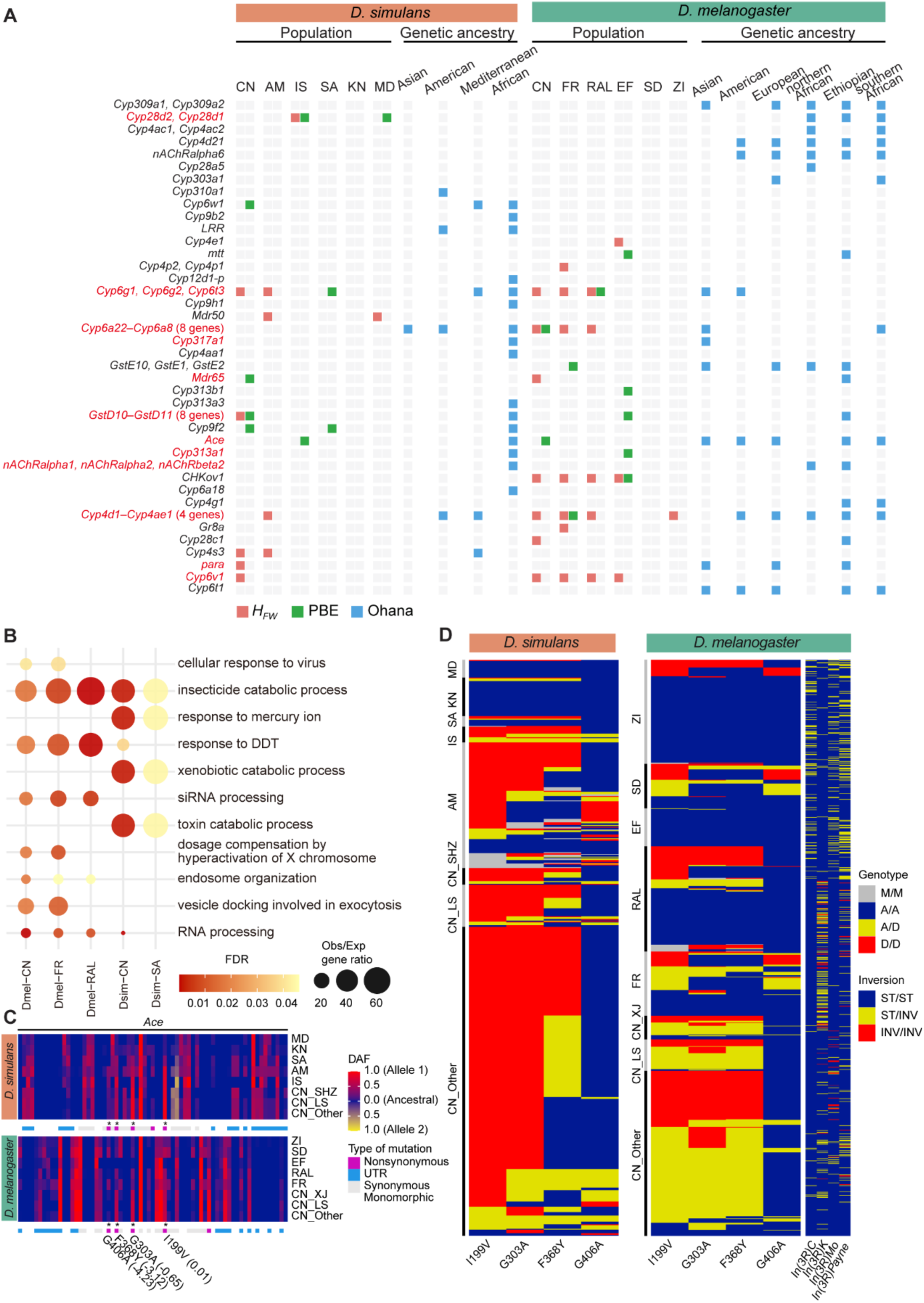
Adaptive convergence and molecular parallelism in insecticide resistance in *D. simulans* and *D. melanogaster*. (A) Selection signals detected by *H*_FW_, PBE, and Ohana for insecticide resistance related genes or gene clusters in major populations or genetic ancestries in *D. simulans* (left) and *D. melanogaster* (right). The names of the genes or gene clusters detected in both species are colored red. (B) Bubble chart of gene ontology (GO) terms with false discovery rates (FDRs) less than 0.05 in at least two populations, showing adaptive convergence within and between species. Each row and column represent a specific GO term and population, respectively. The color and size of the bubble represent the FDR and the observed-to-expected gene ratio of a specific GO term. (C) Heatmap of derived allele frequencies (DAFs) for *Ace* in *D. simulans* (upper) and *D. melanogaster* (lower). Each column represents one of 67 exonic SNPs (MAF > 0.05 in at least one species), and each row corresponds to a population or subpopulation. Derived and ancestral alleles were inferred using the MRCA of the two species. Color gradients represent DAFs, with different hues indicating distinct derived alleles. Missense mutations shared between species (I119V, G303A, F368Y, G406A) are marked with asterisks. Numbers in parentheses denote ESM-scan scores, with negative values indicating predicted deleterious effects. (D) Genotypes of the four shared missense mutations in *Ace* across six major populations of *D. simulans* (left) and *D. melanogaster* (middle). Each column represents a mutation, and each row represents a strain. Genotype color codes: A/A (ancestral homozygous, blue), D/D (derived homozygous, red), A/D (heterozygous, yellow), M/M (missing, gray). Right panel: genotype inference of the 3R inversion in *D. melanogaster*. Inversion status was assigned based on diagnostic SNP frequencies: heterozygous (yellow, 0.1–0.9), standard homozygous (blue, < 0.1), or inverted homozygous (red, > 0.9).

We detected the *Cyp6g1* T316S mutation under positive selection in both species, occurring at higher frequencies in *D. simulans* and intermediate frequencies in *D. melanogaster* (Figure S8C–D). In addition, six nonsynonymous mutations were shared at orthologous sites in three canonical resistance genes—*Cyp6a9, Cyp317a1,* and *Ace*—though allele frequencies differed, with selection signatures usually detected in one species (Figure S11). For example, the *Cyp6a9* G33D mutation segregated at intermediate frequencies in cosmopolitan *D. simulans* but at higher levels in *D. melanogaster*. Similarly, *Cyp317a1* G439N was more frequent in *D. melanogaster* than the orthologous G442N in *D. simulans*. By contrast, *Ace* mutations (I199V, G303A, F368Y, G406A) were generally more frequent in *D. simulans* than in *D. melanogaster* (Figures 5C and S11).

In *D. melanogaster*, the lower frequencies of these *Ace* and *Cyp6g1* variants appear linked to chromosomal inversions (Figure 5D and Table S21). The *Cyp6g1* T316S mutation was significantly associated with *In(2R)Ns* (*P* = 0.015), while *Ace* mutations (I199V, G303A, F368Y) correlated strongly with *In(3R)C*, *In(3R)Mo*, and *In(3R)Payne* but rarely with *In(3R)K* (*P* < 0.001). The *Ace* G406A mutation specifically associated with *In(3R)C* (*P* = 0.001). Thus, chromosomal inversions in *D. melanogaster* likely maintain adaptive mutations in insecticide genes at intermediate frequencies by suppressing recombination, whereas their absence in *D. simulans* facilitates more rapid fixation under positive selection.

## Discussion

The extent to which evolution is repeatable across lineages occupying similar environments remains uncertain. Although natural selection may drive populations toward higher fitness peaks, stochastic forces—including mutation, recombination, drift, and demography—render outcomes inherently unpredictable^88^. Here, we addressed this question by performing a comprehensive analysis of convergent adaptation in two globally distributed sibling species, *D. melanogaster* and *D. simulans*. Using genomic data from over 2,000 strains across continents, we estimate that ∼10% of genes undergo convergent adaptation. This convergence is widespread and functionally coherent at the pathway and gene levels but rare at the nucleotide level, indicating that selection primarily targets functional modules rather than identical sites.

Our results reveal continuous convergence across multiple evolutionary timescales (Figure 6). Divergence-based MKT identified 1,255 positively selected genes in *D. simulans* and 620 in *D. melanogaster*, with 208 (12.5%) shared. Population-level scans (*H*_FW_, PBE) detected 996 selected genes in *D. simulans* and 1,246 in *D. melanogaster*, with 186 (9.1%) shared. Ancestry-aware Ohana analysis identified 611 shared genes (13.3%). Convergent adaptation was notably pronounced among insecticide resistance genes, with selection detected in 64 genes in at least one species and substantial overlap (20 genes, 31.3%; Figure 6). This heightened convergence likely reflects intense selection driven by widespread insecticide use in agriculture. By contrast, among >10,000 selected SNPs per species, only 87 overlapped at orthologous sites. Cage experiments under paraquat-induced oxidative stress further supported this pattern: significant convergence occurred at the gene level, while overlap at nucleotide sites was negligible. Together, these findings highlight a hierarchical pattern—robust convergence at pathways, intermediate convergence at genes, but rare convergence at sites.

**Figure 6.**
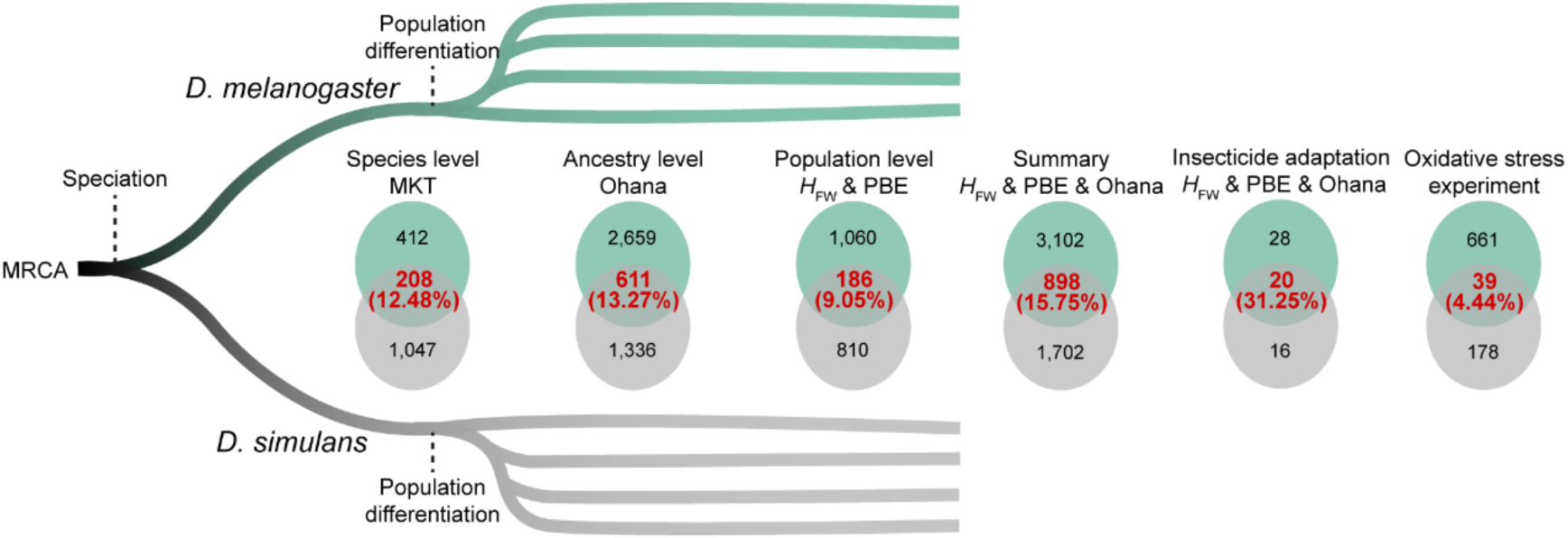
Summary of adaptive convergence in *D. melanogaster* and *D. simulans* across evolutionary timescales. Solid lines depict the evolutionary trajectories of the two species. Venn diagrams illustrate the number and overlap of convergently selected genes identified by different selection detection methods across various timescales. The number and overlap of genes under positive selection related to insecticide resistance or associated with oxidative stress are also highlighted.

We found little evidence for recent interspecific gene flow contributing to convergent adaptation. Specifically, we quantified genomic similarity across 1,335 *D. melanogaster* and 681 *D. simulans* genomes using *k*-mer-based angular similarity via sourmash^89^, which accounts for both presence and frequency of *k*-mers by measuring the angular distance between high-dimensional frequency vectors. In non-repetitive regions (*k* = 31), within-species angular similarity (median: 42.3% in *D. melanogaster*, 48.8% in *D. simulans*) significantly exceeded interspecific similarity (16.6%; Figure S11F). A similar pattern was observed in a subset of 898 adaptively convergent genes, where median similarity within species remained high (49.5– 50.6%) but low between species (17.0%). These trends persisted at *k* = 51, though absolute values decreased slightly (Figure S11G). Thus, our data strongly indicate that convergent adaptation is not driven by recent gene flow; otherwise, adaptive loci would show elevated interspecific similarity compared to background levels.

In summary, our multi-layered framework captures adaptive evolution across lineages and timescales. The results support a model in which shared environmental pressures, such as insecticide exposure, repeatedly channel adaptation toward common phenotypes, yet the precise genetic routes remain lineage-specific. Natural selection thus fosters predictable convergence at functional levels, but stochastic forces and historical contingencies shape the underlying molecular paths.

## Materials and methods

### Genome sequencing

The *D. simulans* strain *w*^501^ (stock number: 14021-0251.195) was obtained from the National *Drosophila* Species Stock Center at Cornell University for reference genome sequencing and assembly. The flies were maintained in the laboratory as random-bred closed stock without further inbreeding prior to sequencing. To generate long reads for genome assembly, we used a combination of third generation sequencing technologies, including Pacific Biosciences high-fidelity sequencing (PacBio HiFi) and Oxford Nanopore Technology (ONT), as well as high-throughput chromosome conformation capture (Hi-C) and Illumina short-read sequencing. For ONT, PacBio HiFi and Hi-C, DNA extraction, library preparation, and sequencing were performed by GrandOmics Biosciences, whereas Illumina short-read sequencing was performed by Annoroad Gene Technology (Beijing).

For nanopore sequencing, genomic DNA was extracted from 300 male flies of the *w*^501^ strain by GrandOmics BAC-long DNA extraction kit (XJZZ-BDE-003). The library was prepared using Ligation Sequencing DNA V14 (SQK-LSK114) according to the guidelines of Oxford Nanopore Technologies. Subsequently, sequencing was performed on the Nanopore PromthION P48 platform. ONT sequencing data in POD5 format were processed using Dorado (https://github.com/nanoporetech/dorado) in “sup” mode for quality control. Low-quality reads were filtered out, resulting in high-quality reads for subsequent analysis. This process yielded approximately 50.31 Gb of sequencing data, corresponding to a depth of approximately 322×.

For PacBio HiFi sequencing, Qiagen Genomic-tips were used for genomic DNA extraction from 300 male flies. After DNA quality control, libraries were prepared using the SMRTbell Prep Kit 3.0. The ligated DNA fragments were purified and recovered, and the size of the target fragments was selected using BluePippin (Sage Science, USA), yielding the final SMRTbell library. The prepared library, containing a defined concentration and volume of DNA templates and enzyme complexes, was loaded onto the PacBio Revio sequencing platform. The automated workstation transferred the library to SMRT cells, initiating single-molecule real-time sequencing. The sequencing data were subjected to quality control using SMRT Link V13.0 with default parameters, where failed reads were filtered out, including low-quality reads with an average quality score (rq < 0.99) and adapter sequences. This process yields approximately 18.76 Gb sequencing data, corresponding to a depth of approximately 120×.

For Hi-C library construction and sequencing, 100 male flies were used for genomic DNA extraction, and the DNA was further digested by *Dpn*II. Short-read sequencing was performed on the MGI DNBSEQ-T7 platform, with a paired-end read length of 150 bp (PE150). This process yields approximately 29.26 Gb of sequencing data, corresponding to a depth of approximately 187×.

For Illumina short-read sequencing, genomic DNA was extracted from 20 male or female flies via a TIANamp Genomic DNA Kit (DP304). A VAHTSTM PCR-Free DNA Library Prep Kit for Illumina (Vazyme, ND602-01) was used for library preparation. The cluster generation and sequencing were performed on the NovaSeq 6000 platform, with a paired-end read length of 150 bp. This process yields approximately 24.21 Gb sequencing data, corresponding to a depth of approximately 155×.

### Genome assembly and polishing

The genome assembly was generated using hifiasm (UL)^90^ version 0.19.5-r587. The PacBio HiFi, ONT, and Hi-C data (Table S1) were simultaneously used as inputs with the hifiasm parameters “--ul”, “--h1”, and “--h2”, respectively, while the other settings were kept as defaults. The draft genome in GFA format was converted to FASTA format using gfatools version 0.4-r214-dirty (https://github.com/lh3/gfatools) with default parameters. The draft genome subsequently underwent additional steps, including filtering, polishing, and duplication purging. To remove DNA contamination, FCS-GX^91^ version 0.4.0 was used to identify contaminants in the draft genome. The “--tax-id” parameter was set to 7240 to specify the species as *D. simulans*, while the other parameters were left as defaults. The database used for contamination detection was downloaded from NCBI (https://ftp.ncbi.nlm.nih.gov/genomes/TOOLS/FCS/database/latest/all.gxs; accessed on September 26, 2023). On the basis of the results from the FCS-GX, two contigs identified as having a non-insect origin were removed from the draft genome (Table S3).

Mitochondrial DNA (mtDNA) sequences were identified from the draft genome using MitoHiFi^92^ version 3.2. The analysis used the *D. simulans* mitochondrial reference genome (TT00 strain, NCBI accession number: NC_005781.1) as the reference, with the parameters set as “-p 90 -o 5” to specify a minimum sequence similarity of 90% and the use of translation table 5 (the invertebrate mitochondrial genetic code). All other settings were left as defaults. On the basis of the MitoHiFi results, the best matching sequence was retained, while 20 redundant mitochondrial sequences were removed.

To remove duplications of the draft genome assembly, purge_dups^93^ version 1.2.5 was used with default parameters, incorporating minimap2^94^ version 2.26-r1175, to identify and remove two redundant sequences. NextPolish2^94^ version 0.2.0 was subsequently used to perform one round of polishing on the duplication- and contamination-free genome using Illumina and PacBio HiFi sequencing data with default parameters. The final genome assembly, after contamination filtering, duplication purging and polishing, consisted of 28 contigs with a total length of 156,078,890 bp, and was used for downstream analyses.

### Quality assessment of the *D. simulans* genome

The quality of the *D. simulans* genome was evaluated from the perspectives of completeness, continuity, and accuracy. Hi-C contact density was quantified using 3D-DNA^95^ version 190716 and Juicer^96^ version 1.6. The contact density between 100-kb windows was visualized using plotHicGenome version 0.1.0 (https://github.com/Atvar2/plotHicGenome). To assess genome completeness, Compleasm version 0.2.4 was used with the parameter “-l diptera” to evaluate the presence of benchmarking universal single-copy orthologs (BUSCOs) from the diptera_odb10 dataset.

The completeness of the genome assembly was further evaluated using metrics such as remapping rates and the breadth and depth of coverage on the basis of the remapping of sequencing data. ONT and PacBio HiFi sequencing reads were mapped to the genome using minimap2, with the parameters “-ax map-ont” and “-ax map-hifi”, respectively, while other settings were left as defaults. Illumina sequencing reads were mapped to the reference genome using BWA-MEM version 0.7.17-r1188 with the “-M” parameter to mark shorter split hits as secondary, whereas all other settings were set to defaults. Alignments with a mapping quality less than 20 were filtered out via SAMtools. The average coverage depth across 100-kilobase (kb) sliding windows was calculated via SAMtools coverage with the parameter “-mean”.

Additionally, as recommended by Genome Continuity Inspector^97^ (GCI) version 1.0, ONT and PacBio HiFi reads were mapped to the reference genome using Winnowmap version 2.03 with the parameters “-x map-ont” and “-x map-pb”, respectively, while other settings remained default. The results from minimap2 and Winnowmap were processed with GCI using default settings to detect potential assembly issues and quantify genome continuity. To quantify the error rate, *k*-mer databases (*k* = 19) were constructed using PacBio HiFi or Illumina sequencing data with Meryl^98^ version 1.3, and the quality value (QV) and error rate were then evaluated using Merqury^98^ version 1.3.

To compare the syntenic relationships between the reference genome and other genome versions, our genome assembly was analyzed alongside the *D. melanogaster* reference genome^99^ (dmel_r6.54, NCBI accession: GCF_000001215.4) and previously published *D. simulans* genomes^37^ (Dsim3.1, NCBI accession: GCF_016746395.2; dsim_r2.02, NCBI accession: GCF_000754195.3) using NGenomeSyn version 1.41. For the analysis, scaffolds smaller than 1 Mb were excluded by setting the parameters “-MinLenA 1000000 -MinLenB 1000000”. Minimap2 was used for alignment with the parameter “-MappingBin minimap2”, while other settings were kept as defaults.

### Identification of Y-linked contigs

To identify Y-linked contigs, we obtained Illumina paired-end (PE150) reads from male and virgin female PCR-free genomic libraries (Table S1). These reads were mapped separately to the reference genome using BWA-MEM, and reads with mapping quality or base quality below 10 and multi-mapped reads were excluded. We subsequently used SAMtools to calculate the depth of coverage per site and summarized the depth across each contig. Contigs with a female-to-male depth ratio less than 0.2 were considered Y-linked. To further validate the Y-linked contigs, we excluded those without syntenic blocks corresponding to previously identified Y-linked contigs^100^, resulting in a total of 12.64 Mb of Y-linked sequences.

### Gene annotation

To annotate protein-coding genes (PCGs) in the *D. simulans* genome, we integrated evidence from protein homology, RNA sequencing (RNA-seq), full-length transcriptome data (ISO-seq), and *ab initio* predictions (Figure S12).

Prior to PCG annotation, repetitive sequences were annotated using four approaches: RepeatMasker (https://github.com/Dfam-consortium/RepeatMasker), RepeatModeler^101^, Extensive de-novo TE Annotator^102^ (EDTA), and tandem repeats finder^103^ (TRF). RepeatMasker version 4.1.1 utilized RepBase version 20181026 as a reference database to identify homologous repeat sequences with the parameter “-species 7240”, defining the species as *D. simulans*, while other settings were kept as defaults. RepeatModeler version 2.0.1 and EDTA version 1.9.6 were executed with default parameters. TRF version 4.09.1 was used to predict tandem repeat sequences, with the parameters set to “2 7 7 80 10 50 500 -f -d -m -h”. The identified repeat sequences from these methods were classified and summarized to generate a nonredundant repeat sequence coordinates. These coordinates were used to soft-mask the genome with bedtools maskfasta version 2.30.0 using the “-soft” parameter. The soft-masked genome was then used for PCG annotation.

For homology-based predictions, protein sequences were analyzed via Exonerate^104^, Miniprot^105^, and GeMoMa^106^. Exonerate utilized protein sequences from 12 *Drosophila* species (*D. ananassae* r1.3, *D. erecta* r1.3, *D. grimshawi* r1.3, *D. melanogaster* r6.54, *D. mojavensis* r1.3, *D. persimilis* r1.3, *D. pseudoobscura* r3.2, *D. sechellia* r1.3, *D. simulans* r2.02, *D. virilis* r1.2, *D. willistoni* r1.3, and *D. yakuba* r1.3), downloaded from FlyBase (http://ftp.flybase.net/genomes/; accessed on December 1, 2023). Only the longest protein sequence for each gene was retained. Initially, the protein sequences were aligned to the soft-masked genome using tblastn version 2.15.0 to identify approximate locations of the proteins in the genome. The parameters were set to “-outfmt 6 -evalue 1e-5 -soft_masking true”, with all other settings set to defaults. The precise alignment of proteins to their corresponding genomic regions was subsequently performed using exonerate (version 2.4.0) with the parameters “--percent 20 --score 0 --model protein2genome --querytype protein --targettype dna --showvulgar no --softmasktarget yes --showalignment no --showtargetgff yes --showcigar no --bestn 1”. The output GFF file from this process constituted gene set 1.

Miniprot version 0.12-r237 was used to analyze the 12 *Drosophila* (mentioned above) genomes as homologous references for gene prediction. The parameter “-j 2 --gff” was used to set the splicing mode to mammalian, which is also compatible with *Drosophila*^105^. All other parameters were set to default values. The resulting GFF file from this analysis constituted gene set 2.

GeMoMa version 1.7.1 was utilized for homology-based gene prediction by incorporating genomic and annotation files from both *D. melanogaster* and *D. simulans*, as well as RNA-seq data as additional evidence (Table S22). The raw RNA-seq data were retrieved from SRA, NCBI (accessed on August 29, 2023). Quality control (QC) of the RNA-seq data was performed using fastp version 0.20.1 with the parameters “-u 50 -n 5 -q 20 -5 -3 -W 4 -l 50”. This process removed reads where more than 50% of the bases were of a quality less than 20, reads containing more than 5 unknown bases (N), and reads shorter than 50 bp. Additionally, 4 bases were trimmed from both the 5′ and 3′ ends if their quality scores were below 20, and Illumina adapter sequences were removed. All other parameters were set to their defaults. The QC-filtered RNA-seq data were aligned to the soft-masked reference genome using HISAT2^107^ version 2.2.1 with default parameters. The RNA-seq alignment BAM files, along with the genomic and annotation files from *D. melanogaster* and *D. simulans*, were input into GeMoMa for gene prediction. GeMoMa was configured with the parameters “AnnotationFinalizer.r=NO p=false o=true tblastn=true AnnotationFinalizer.u=YES r=MAPPED”, while all other parameters were set to defaults. The resulting GFF file from this analysis constituted gene set 3.

To predict PCGs using RNA-seq evidence, Mikado^108^ software version 2.3.4 was used with the daijin workflow. The workflow was executed with the parameters “-al hisat -as stringtie trinity -m permissive”. Protein sequences from arthropods in OrthoDB^109^ version 10 and RNA-seq data (Table S22) were incorporated as evidence. The daijin workflow used the following tools: HISAT2^107^ version 2.2.1 for RNA-seq read alignment, SAMtools^110^ version 1.12 for BAM file manipulation, Trinity^111^ version 2.9.1 and StringTie^112^ version 2.2.1 for de novo transcriptome assembly, Portcullis^113^ version 1.2.4 for splice junction filtering, Mikado^108^ version 2.3.4 for selecting the best transcript models, GMAP^114^ version 2023-10-10 for transcript alignment, and DIAMOND^115^ version 2.1.8 for protein homology searches. The GFF file generated by this workflow was designated gene set 4.

The prediction of PCGs was supported by full-length transcriptome (ISO-seq) data, processed using the nf-core/isoseq workflow^116,117^ version 1.1.5-g1d5244a and PASA^118^. Raw ISO-seq data in the form of subread BAM files were downloaded from NCBI (Table S22). These files were processed with the nf-core/isoseq pipeline (with Nextflow^119^ version 23.10.0) using the parameter “aligner: ’minimap2’” for assembling full-length transcripts on the basis of primer sequences. The pipeline integrates multiple tools, including pbccs version 6.4.0 (https://github.com/PacificBiosciences/ccs.git), bamtools^120^ version 2.5.2, lima version 2.7.1 (https://github.com/PacificBiosciences/barcoding.git), gs-tama^121^ version 1.03, and isoseq3 version 3.8.2 (https://github.com/PacificBiosciences/IsoSeq.git). These tools were utilized to identify mRNA sequences on the basis of primer sequence positions, remove poly(A) tails and convert the sequences to FASTA format, resulting in a dataset of full-length transcripts. Using the full-length transcript sequences, the annotations of the PCGs were further refined with both nf-core/isoseq and PASA. The nf-core/isoseq workflow used minimap2 version 2.24-r1122 and tama to map the transcripts onto the genome. The resulting BED12 file represents gene set 5.

Full-length transcriptome sequences were input into PASA^118^ version 2.5.3 for gene annotation, using the “Launch_PASA_pipeline.pl” workflow with the parameters set to “-C -R --ALIGNERS gmap --TRANSDECODER --MIN_AVG_PER_ID=60 --MIN_PERCENT_ALIGNED=60”. This workflow invoked GMAP version 2023-10-10 and transdecoder version 5.5.0 (https://github.com/TransDecoder/TransDecoder.git). The resulting GFF file represents gene set 6.

*Ab initio* prediction of PCGs was performed using BRAKER3^122–124^ version 3.03, with protein sequences from arthropods in orthoDB^109^ version 10 used as input evidence. The workflow utilized ProtHint^125^ version 2.6.0, GeneMark-EP+^125^ version 1.0.0, DIAMOND^115^ version 2.1.8, Spaln^126^ version 3.0.1, and Augustus^127,128^ version 3.4.0 for gene prediction, with default parameters. The resulting GFF file corresponds to gene set 7.

To integrate the four types of evidence, the study utilized EVidenceModeler (EVM)^129^ version 2.1.0 to combine the aforementioned seven gene sets into a nonredundant PCG annotation file. The weights for protein evidence, RNA sequencing evidence, full-length transcriptome evidence, and de novo predictions were set to 8, 10, 10, and 1, respectively. Default parameters were used. In total, 15,336 nonredundant PCGs were annotated. The annotation file was subsequently updated using PASA, which used full-length transcript sequences and transcripts assembled by Mikado. This process refined the annotations of neighboring and overlapping genes, transcripts, and untranslated regions (UTRs), resulting in a final annotation of 15,522 PCGs.

### Functional annotation of genes

The annotated PCGs may contain transposable element (TE) genes and genes without functional domains. Therefore, three methods were employed for the first round of functional annotation of the protein sequences of the PCGs: eggNOG-mapper^130^ version 2.1.12, InterProScan^131^ version 5.59_91.0, and BLASTP.

eggNOG-mapper was accessed via the web interface (http://eggnog-mapper.embl.de/) using the eggNOG 5.0 database^132^. The parameters were set to “-m diamond --evalue 0.001 –score 60 --pident 40 --query_cover 20 --subject_cover 20 --itype proteins --tax_scope auto --target_orthologs all --go_evidence non-electronic --pfam_realign none --report_orthologs”. InterProScan utilizes several databases, including AntiFam 7.0, CDD 3.18, Coils 2.2.1, FunFam 4.3.0, Gene3D 4.3.0, Hamap 2021_04, MobiDBLite 2.0, PANTHER 17.0, Pfam 35.0, PIRSF 3.10, PIRSR 2021_05, PRINTS 42.0, ProSitePatterns 2022_01, ProSiteProfiles 2022_01, SFLD 4, SMART 7.1, SUPERFAMILY 1.75, and TIGRFAM 15.0. The parameters were set to “-iprlookup -pa -dp -goterms”. BLASTP was performed using DIAMOND version 2.1.8 for sequence alignment against the SwissProt database (downloaded from https://ftp.ncbi.nlm.nih.gov/blast/db/FASTA/swissprot.gz on January 2, 2024) via the parameters “--evalue 1e-5 --max-target-seqs 5”. Additionally, TEsorter^133^ version 1.4.6 was used to annotate the TE proteins, with the parameters set to “-db rexdb-metazoa -eval 1e-6”. Combining the four annotation methods, genes associated with TE-related proteins were removed from the annotation file. This resulted in the retention of 15,131 annotated PCGs.

A second round of functional annotation was performed. The methods for eggNOG-mapper and InterProScan remained the same as those used in the first round. Additionally, BLASTP was used to align protein sequences against three databases: the SwissProt database, the nr database (downloaded from https://ftp.ncbi.nlm.nih.gov/blast/db/FASTA/nr.gz on January 2, 2024), and the protein sequences of *D. melanogaster* and *D. simulans* (downloaded from FlyBase https://ftp.flybase.net/genomes/ on December 1, 2023).

Combining the results from these five methods, genes associated with TEs were further removed, and only those genes annotated with functional domains or closely related proteins by at least one method were retained. In total, 14,927 PCGs were included in the final annotation file.

In addition, noncoding RNAs were annotated using Infernal version 1.1.4 and tRNAScan-SE^134^ version 2.0.9. Infernal utilized the Rfam database version 14.10, which was downloaded from EBI (https://ftp.ebi.ac.uk/pub/databases/Rfam/14.10/, accessed on December 4, 2023). Default parameters were used for both tools. We further identified miRNAs basis of their similarity with previously reported miRNA sequences from miRbase^135^ Release 22.1 (https://www.mirbase.org/download/hairpin.fa, accessed Jan. 12, 2025). Each miRNA primary transcript from 12 *Drosophila* species (*D. melanogaster*, *D. pseudoobscura*, *D. sechellia*, *D. simulans*, *D. yakuba*, *D. erecta*, *D. ananassae*, *D. persimilis*, *D. willistoni*, *D. mojavensis*, *D. virilis*, and *D. grimshawi*) was mapped to the reference genome of *D. simulans* using BWA-MEM with the parameter “-M”, while the mapped miRNAs with mapping quality below 20 and soft or hard clipping were filtered out. Genomic regions without mismatches or indels compared with those of miRbase were considered miRNAs, whereas those with no more than five mismatches or indels were considered putative miRNAs. In total, 271 miRNAs were identified in the genome of *D. simulans*, with 165 perfectly matched miRNAs and 155 putative miRNAs identified by miRbase, and 11 additional miRNAs identified only by Infernal.

The neutral sites in the *D. simulans* genome consist of two parts: short introns and fourfold degenerate (4d) sites within coding regions. The annotation of the 4d sites was based on the annotation GFF file. A database was generated using SnpEff^136^ version 5.0, and three possible mutations for each site were annotated; only sites where all three alleles corresponded to synonymous mutations were retained. Additionally, sites at 8–30 bp of introns with a length of 45–60 bp were also considered neutral^62^, whereas sites with any overlap with UTRs or exons from other genes were removed.

For *D. melanogaster*, the genome annotation file was downloaded from FlyBase (version r6.54). This file was then used to annotate the neutral sites in the *D. melanogaster* genome, following the same method used for *D. simulans*.

### Genome coordinate conversion, genomic divergence and ancestral allele inference

To convert the genome coordinates between *D. melanogaster* and *D. simulans*, we used LastZ^137^ version 1.04.22 for sequence alignment. The kentUtils pipeline, specifically the scripts “doBlastzChainNet.pl” version 1.33 and “doRecipBest.pl” version 1.12, was used to map the *D. simulans* genome to those of *D. melanogaster* and the reciprocal best alignments were retaining. The parameters were set according to the UCSC 124-way multiple alignments for *D. melanogaster*, with the following configuration: “M=0 K=3000 Y=3400 L=4000 E=30 H=2000 O=400 T=1”, and the HoxD55 scoring matrix was used (see http://genomewiki.ucsc.edu/index.php?title=Dm6_124-way_conservation_lastz_parameters). LastZ was also used for alignments between different versions of the *D. simulans* genome (Dsim2 and Dsim3), and between *D. simulans* and the *D. sechellia* genome version r1.3. Additionally, chain files for the alignment of *D. melanogaster* to other species (*D. simulans* DroSim2, *D. yakuba* DroYak3) were downloaded from UCSC (https://hgdownload.soe.ucsc.edu/goldenPath/dm6/; accessed July 1, 2021). Reciprocal best chain files were then used to convert *D. melanogaster* or *D. simulans* genome coordinates to those of other species using UCSC LiftOver (https://hgdownload.cse.ucsc.edu/admin/exe/) version 377. We calculated the genomic divergence between the *D. melanogaster* reference genome (Dmel6) and *D. simulans* genomes assembled in this study and in previous studies (Dsim2 and Dsim3) using the alignments described above.

Ancestral allele states at each site were estimated using est-sfs^138^ version 2.03. To infer the ancestral alleles of *D. simulans*, polymorphism data from *D. simulans*, and the genotypes of *D. sechellia* and *D. melanogaster* were used. On the basis of the phylogeny, *D. melanogaster* was considered the outgroup, and this analysis allowed the estimation of alleles at the most recent common ancestor (MRCA) of *D. simulans* and *D. sechellia*. Similarly, to estimate the ancestral alleles of *D. melanogaster*, polymorphism data from *D. melanogaster*, and the genotypes of *D. simulans* and *D. yakuba* were used to infer alleles at the MRCA of *D. melanogaster* and *D. simulans*. The est-sfs used the “Rate-6” model^138^ for these calculations and retained sites where the probability of the ancestral allele state was greater than 0.9.

### Sampling and establishment of iso-female lines

In 2017 and 2020, *D. melanogaster* and *D. simulans* were collected from 58 locations across 22 provinces in China. Both species were collected at 46 locations, whereas at 6 locations, only *D. simulans* or *D. melanogaster* was found. For each female fruit fly collected in the wild, all offspring were used to establish an isofemale line in the laboratory.

### Genome resequencing and variant calling

We conducted whole-genome resequencing on 419 *D. simulans* isofemale lines and 108 *D. melanogaster* isofemale lines. The genomic DNA of 419 *D. simulans* and 108 *D. melanogaster* strains (sim1–sim184, mel111 and mel141 were extracted by the authors, and other samples were extracted by Annoroad) was extracted using the TIANamp Genomic DNA Kit (DP304), with 16–24 male flies used for each isofemale line. All libraries were prepared according to the Watchmaker DNA Library Prep Kit with Fragmentation (7K0019-096) by Annoroad Gene Technology (Beijing), and sequenced using the Illumina NovaSeq 6000 platform with a paired-end read length of 150 bp (PE150) by Annoroad, generating approximately 5 Gb of raw sequencing data for each strain. Combined with previously published data from China^38^, this study obtained whole-genome resequencing data from 419 *D. simulans* strains and 400 *D. melanogaster* strains from 56 locations across China.

A total of 558 *D. simulans* genomes were downloaded from the NCBI SRA using the SRA Toolkit version 3.0.8 (https://github.com/ncbi/sra-tools.git). These genomes were collected from Madagascar, Kenya, Senegal, Equatorial Guinea, South Africa, Israel, Spain, and the United States^52–54,56–58,139–141^. For *D. melanogaster*, the genomic data used in previous studies^38^ were obtained, with 1,430 *D. melanogaster* genomes downloaded for analysis. For both previously published data and data generated in this study, quality assessment of fastq files was performed using FastQC version 0.11.5 (https://github.com/s-andrews/FastQC.git) and multiqc^142^ version 1.9. Quality control was subsequently conducted with fastp version 0.20.1 using the parameters “-w 20 -u 50 -n 5 -q 20 -5 -3 -W 4 -l 50”, which was the same as those used for RNA-seq read processing.

High-quality reads for each strain were mapped to the reference genome using BWA-MEM version 0.7.17-r1188, with the parameter “-M”, while the other parameters were set to defaults. The reference genome for *D. simulans* was the assembly generated in this study, whereas the reference genome for *D. melanogaster* was version 6 downloaded from FlyBase. To remove optical duplicates, Picard version 2.23.9 (https://github.com/broadinstitute/picard.git) was used with the parameter “REMOVE_SEQUENCING_DUPLICATES=true”, leaving other settings as defaults.

We performed joint variant calling on 997 *D. simulans* genomes and 1,547 *D. melanogaster* genomes (Table S9) using FreeBayes version 0.9.21. The parameters used were “--haplotype-length 0 --use-best-n-alleles 4 -m 20 -q 20”. These settings limit consideration to no more than four alleles, a haplotype length of 0, and a minimum read mapping and base quality of 20.

### Variant filtering and annotation

To filter low-quality variants, the Variant Call Format (VCF) files were processed using vcflib^143^ version 1.0.2 and VCFtools^144^ version 0.1.16 with the following criteria: (1) mutations with a quality score (QUAL) below 20 and alternative allele observations (AO) fewer than 5 were removed; (2) genotypes with a sequencing depth of less than 3 were marked as missing; (3) multinucleotide polymorphisms (MNPs) and complex mutations were decomposed into single nucleotide polymorphisms (SNPs) and insertions or deletions (indels); (4) sites within repetitive regions were excluded (annotated by RepeatMasker or UCSC Genome Browser in *D. melanogaster*, and by four aforementioned methods in *D. simulans*); (5) sites with more than 10% missing genotypes were removed; and (6) only biallelic sites were retained. To minimize alignment errors around indels, SNPs located within 3 bp of an indel were excluded. Strains with more than 20% missing data and outliers identified via principal component analysis (PCA) were also removed (described in detail below), leaving a final dataset of 681 *D. simulans* genomes. For *D. melanogaster*, similar sites and strain filtering steps as those used for *D. simulans* were applied, and we included only 22 major populations described in previous studies^38^, retaining 1,335 genomes after filtering. The functional consequences of these SNPs were annotated using SnpEff and an annotation file from FlyBase (version 6.54) for *D. melanogaster* and an annotation file generated in the present study for *D. simulans*.

### Inversion genotyping of *D. melanogaster*

The chromosomal inversion genotypes of *D. melanogaster* were determined using diagnostic SNPs of inversions^145^. For each strain, the average allele frequency of diagnostic SNPs was calculated; those with an average frequency less than 0.1 or greater than 0.9 were considered homozygotes of standard or inverted chromosomes, whereas those with an average frequency 0.1 and 0.9 were considered heterozygotes. Using this method, the genotypes of seven common chromosomal inversions in 108 newly sequenced *D. melanogaster* genomes were identified, including *In(2L)t*, *In(2R)NS*, *In(3L)P*, *In(3R)C*, *In(3R)K*, *In(3R)Mo*, and *In(3R)Payne*, whereas the inversion genotypes of other strains were derived from a previous study^38^.

### Principal component analysis and population structure

Principal component analysis (PCA) and population structure analysis were conducted using neutral sites (including the short introns and 4d sites mentioned above) after filtering out sites with a MAF less than 0.01. The number of sites used in the analysis was 273,514 (chr2: 113,121, chr3: 135,122, chr4: 186, chrX: 25,085) and 213,284 (2L: 48,876, 2R: 44,430, 3L: 41,594, 3R: 47,070, X: 31,314) for *D. simulans* and *D. melanogaster*, respectively. The analyses were conducted based on each chromosome (arm) for *D. simulans* and *D. melanogaster*, while autosomes were also combined into a single dataset for analysis in *D. simulans*. Strains of *D. melanogaster* carrying chromosomal inversions were excluded from the analyses of each chromosome, whereas all strains of *D. simulans* were included. VCF files were converted into PLINK PED and BED formats using VCFtools and PLINK^146^ version 1.90b6.18.

PCA was performed with the smartpca module of EIGENSOFT version 7.2.1 using default parameters. For PCA outlier removal in the strain filtering part, outliers were manually detected and removed according to the PCA results based on both autosomes and the X chromosome. ADMIXTURE^147^ version 1.3.0 was used to infer the genetic component of each strain. The analysis was conducted with the number of genetic ancestry (*K*) ranging from 1 to 10, with 10 replicates for each *K* using random seeds (Table S13). The optimal value of *K* was determined on the basis of the lowest cross-validation error (Table S13). The results for different *K* values were summarized using pong^148^ version 1.4.9 with the parameters set to “--greedy -s.95”, which grouped results with a similarity above 95% into a single model or processed using a greedy algorithm.

### Genetic diversity and population differentiation

The genetic diversity (π) of each population was calculated using pixy^149^ version 1.2.4.beta1 on the basis of all polymorphic sites and strains of 11 and 22 populations of *D. simulans* and *D. melanogaster*, respectively. As Pixy requires an all-site VCF file, a custom VCF file was prepared by filling all non-SNP sites with reference homozygous genotypes (0/0). π was calculated using 200-kb nonoverlapping windows, with the weighted average π determined by dividing the sum of the observed differences (count_diffs) by the sum of the comparisons (count_comparisons). For autosomal π (π_A_), the weighted average was computed as the sum across chromosomes 2, 3, and 4. To account for differences in effective population size (*N_e_*), a corrected X chromosome diversity (π_Xc_) was also calculated as π_X_ multiplied by 4/3.

Additionally, π was calculated for various genomic functional categories, including coding sequences (CDS), 5′ UTRs, 3′ UTRs, introns, and intergenic regions. For these calculations, VCF files containing all sites within each functional category were extracted from the all-site VCF file according to the genome annotation (GFF) files. Overlapping regions between functional categories were removed. Weighted averages for π were calculated using the same method used for the whole genome.

Population differentiation was quantified via the fixation index (*F*_ST_) and absolute nucleotide divergence (*D*_XY_). The pairwise *F*_ST_ was calculated using Weir and Cockerham’s method via VCFtools, whereas *D*_XY_ was computed via Pixy. Pairwise comparisons were performed across 11 and 22 populations of *D. simulans* and *D. melanogaster*, respectively. These analyses included all strains and all sites, with separate calculations for autosomes and the X chromosome. Weighted averages of *D*_XY_ were computed in the same way as for π.

### Phylogenetic analysis and estimation of gene flow among populations

Phylogenetic analysis was performed using TreeMix^64^ version 1.13 on the basis of allele frequency data from nine major *D. simulans* populations (with more than 10 genomes) and 22 *D. melanogaster* populations. Neutral biallelic SNPs with a MAF > 0.01 were used in this analysis. Strains of *D. melanogaster* carrying chromosomal inversions were excluded from the analyses of each chromosome, whereas all strains of *D. simulans* were included. The analyses were conducted separately for autosomes (combining chr2, chr3 and chr4) and chrX for *D. simulans*, and on the basis of 2R, 3L and X for *D. melanogaster*, as all strains of GH and CO contain inversions on 2L or 3R.

Maximum likelihood (ML) trees were constructed to model different numbers of gene flow events. Analyses were conducted using the parameters “-k 50 -root MD/ZI -bootstrap”. This setting uses blocks of 50 SNPs, with the MD population as the root for *D. simulans* and the ZI population as the root for *D. melanogaster*. The number of migration events (*m*) was tested from 0 to 10. For each value of *m*, 100 independent runs were performed using random seeds. The optimal value of m was determined using the second-order rate of change in likelihood (Δ*m*) calculated by the R package OptM version 0.1.6. The result with the highest likelihood for the optimal *m* was selected as the representative result. Visualization of the phylogenetic tree and migration events was performed using plotting_funcs.R script of TreeMix.

We subsequently used ADMIXTOOLS2^150^ version 2.0.0 to calculate the *f*-statistics for these nine *D. simulans* populations and constructed an admixture graph. In this analysis, *D. sechellia* was used as the outgroup. Neutral biallelic SNPs with MAF > 0.01 were used and analyses of autosomes (combining chr2, chr3, and chr4) and the X chromosome were performed separately. The *f_2_*-statistics were computed for all populations with the extract_f2 function, using the parameter “blgsize = 4000” to set the block size of the SNP to 4000 bp. The optimal graph topologies were subsequently searched by the find_graphs function on the basis of the *f* statistics. We considered admixture events ranging from 0 to 3, and 20 independent runs were performed for each specified number of admixture events. The search process started with a random topology for each run. Within each run, the topology was iteratively optimized for a maximum of 200 iterations; optimization was terminated if the likelihood score did not improve after 20 consecutive iterations. The model with the lowest score in 20 runs was selected as optimal for each specified number of admixture events. Finally, the optimal graph topology was accessed using the compare_fits function with the parameter “nboot = 100”, which compared out-of-sample scores and graph fitness between graphs with different numbers of admixture events on the basis of bootstrapped SNP blocks. A graph with a greater number of admixture events was selected only if it exhibited a significantly lower score (*P* < 0.01).

We used Dsuite^151^ version 0.4r43 to calculate Patterson’s *D*^66^ statistic for different trios of both *D. simulans* and *D. melanogaster* populations. *D. sechellia* was used as the outgroup for *D. simulans* populations, whereas *D. simulans* was used as the outgroup for *D. melanogaster* populations. We used all SNPs and all strains in this analysis, whereas chromosome arms containing inversions were excluded in *D. melanogaster*. We included only 46 trios for *D. simulans* populations and 1,066 trios for *D. melanogaster* populations that had consistent topologies inferred by TreeMix and had consistent topologies based on both autosomes and the X chromosome (Table S14). To identify significant gene flow within a trio, we applied the same criteria as those used in a previous study^38^ as follows: *D* > 0.05, *z* score > 3, and FDR < 0.05.

### Demographic history of *D. simulans*

In the following analysis of *D. simulans*, mutation rates were set to 4.51 × 10^-9^ per site per generation^56^, and a generation time of 15 generations per year was applied, following a previous study^152^ based on a North American population of *D. melanogaster*. Effective population size (*N_e_*) dynamics were reconstructed using Stairway Plot 2 version 2.1.1 on the basis of the unfolded site frequency spectrum (uSFS). Neutral loci from all strains in representative populations (MD, IS, AM, and CN) were used for this analysis, with one allele randomly sampled from each genome to minimize the effects of inbreeding. The uSFS for each population was generated using ∂a∂i^153^ version 2.1.1 with the “projections” parameter set to the number of samples that maximized diversity, thereby mitigating the impact of missing genotypes.

We used the Python package moments^154^ version 1.1.10 to perform model fitting and parameter estimation for several trios: ((IS, KN), MD), ((IS, SP), MD), ((AM, IS), MD), ((CN_LS, IS), MD), ((CN_Other, CN_LS), IS), and ((CN_Other, CN_SHZ), IS). The demographic models we used were similar to those used in the previous study^38^. For ((IS, KN), MD), ((IS, SP), MD), ((AM, IS), MD) and ((CN_LS, IS), MD), we compared four demographic models: Model 1 (OOA_SyM) assumes symmetric migrations and constant population sizes; Model 2 (OOA_AsyM) assumes asymmetrical migrations and constant population sizes; Model 3 (OOA_BG_SyM) assumes symmetric migrations, constant population sizes for MD, and population size changes in other populations; Model 4 (OOA_BG_AsyM) is similar to Model 3 but allows for asymmetrical migrations (Figure S13, Models 1–4). For ((CN_Other, CN_LS), IS) and ((CN_Other, CN_SHZ), IS), we compared six demographic models: Model 5 (COS_SyM) assumes an ancient bottleneck, symmetric migration and constant population sizes; Model 6 (COS_AsyM) is similar to Model 5 but allows for asymmetrical migrations; Model 7 (COS_BG_SyM) and Model 8 (COS_BG_AsyM) are similar to Model 5 and Model 6, respectively, but allow for population size changes after the divergence of different populations; Model 9 (COS_BG2_SyM) and Model 10 (COS_BG2_AsyM) are similar to Model 7 and Model 8, respectively, but allow for population growth during the ancient bottleneck. The three-dimensional joint folded site frequency spectrum (SFS) for each trio was generated using neutral biallelic SNPs from autosomes by the Spectrum.from_data_dict function in ∂a∂i. To reduce the computational cost, populations with more than 20 samples were projected down to 20 when generating the SFS. The joint SFS was subsequently used to fit these models using moments_pipeline (https://github.com/dportik/moments_pipeline.git). In this pipeline, four rounds of model optimization (replicates = 200, 200, 200, 200; maxiter = 100, 100, 100, 100; fold = 3, 2, 2, 1) with the Nelder–Mead method (optimize_log_fmin) were conducted, and the log-likelihood was inferred by a multinomial approach. We selected the replicate with the highest log-likelihood for each model and then calculated Akaike weights and chose the model with the highest Akaike weight as the best-fitting model for each trio. Details about the model selection and estimated parameters are provided in Table S16. Confidence intervals (CIs) were inferred by generating 100 simulated datasets using msms version 3.2rc, re-estimating parameters under the best-fitting model, and calculating the 2.5–97.5% percentiles for each parameter with the re-estimated results. In the simulation, we set the mutation rate to 4.51 × 10^-9^ per site per generation and the recombination rate to 1 cM/Mb.

### McDonald–Kreitman test

To investigate long-term adaptive protein evolution in *D. simulans* and *D. melanogaster*, the McDonald–Kreitman test (MKT) was conducted on coding regions on the basis of 681 *D. simulans* and 1,335 *D. melanogaster* genomes (Figure S5A). The ancestral alleles at each site were inferred as described previously. To increase comparability between the two species, the alleles at the most recent common ancestor (MRCA) node of *D. simulans* and *D. melanogaster* were considered the ancestral state. Sites were categorized into three groups on the basis of their derived allele frequency (DAF): (1) sites with a DAF exceeding 0.95 were considered fixed substitutions within the species; (2) sites with a DAF between 0.05 and 0.95 were regarded as polymorphic; and (3) sites with a DAF below 0.05 were considered monomorphic. On the basis of the effects of each mutation on amino acid coding, the sites in each species were further classified into four categories: synonymous substitution (*D*_s_), synonymous polymorphism (*P*_s_), nonsynonymous substitution (*D*_n_), and nonsynonymous polymorphism (*P*_n_). Following previous studies^14,69^, a minimum threshold of 6 mutations per gene was applied for each category (polymorphism, substitution, synonymous, and nonsynonymous) for Fisher’s exact test. The Benjamini–Hochberg procedure was used to control the false discovery rate (FDR). The values of *α* and the direction of selection (DoS) were calculated using established formulas^155,156^. Genes with *P* < 0.05 were considered to significantly deviate from neutral expectations, and those with both *α* and DoS values greater than 0 were considered to exhibit signatures of positive selection.

### Fay and Wu’s *H*

Fay and Wu’s *H* (*H*_FW_) was calculated for six representative populations of *D. simulans* (CN, IS, AM, SA, KN, and MD) and *D. melanogaster* (CN, RAL, FR, SD, EF, and ZI). These populations are geographically comparable between species, with populations from ancestral ranges (MD for *D. simulans*, ZI for *D. melanogaster*), non-ancestral sub-Saharan African regions (SA and KN for *D. simulans*, SD and EF for *D. melanogaster*), America (AM for *D. simulans*, RAL for *D. melanogaster*), Mediterranean regions (IS for *D. simulans*, FR for *D. melanogaster*) and Asia (CN for both species). All sites with inferred ancestral alleles and all strains were included in the analysis. Initially, VCF files were converted to diploid HapMap format using TASSEL^157^ version 5.2.40-3, with the inferred ancestral states manually added to the output. On the basis of this file, *H*_FW_ was calculated for 12 populations independently using Variscan^158^ version 2.0 with a sliding window size of 4 kb and a step size of 1 kb. For each 4-kb window, *H*_FW_ values were averaged on the basis of the total number of segregating sites. Thresholds were defined as the top 1% of windows with the lowest *H*_FW_ values for each species (*D. simulans*: *H*_FW_ < −0.28; *D. melanogaster*: *H*_FW_ < −0.53). Windows with an average *H*_FW_ lower than the threshold and containing no less than 50 segregating sites were considered candidate windows. Polymorphic sites within these windows with derived allele frequencies (DAFs) greater than 0.2 were considered to be under selection, and the corresponding genes were identified as candidate targets of selection.

### Population branch excess

To detect population-specific natural selection on the basis of population differentiation, population branch excess (PBE) was calculated using the fixation index (*F*_ST_). The analysis was conducted across six major populations of *D. simulans* (CN, IS, AM, SA, KN, and MD) and *D. melanogaster* (CN, RAL, FR, SD, EF, and ZI), as well as within subpopulations within CN (CN_SHZ, CN_LS, and CN_Other for *D. simulans* and CN_XJ, CN_LS, and CN_Other for *D. melanogaster*). These populations were grouped into cosmopolitan populations, sub-Saharan African populations, and CN subpopulations.

Following established methodologies^159^, outgroup populations were included in each trio to identify branch-specific signals of selection. A total of three tests were performed for each species. For *D. simulans*, the following phylogenetic subtrees were used: (((CN, AM), IS), SA), (((KN, MD), SA), CN), and (((CN_Other, CN_LS), CN_SHZ), IS). For *D. melanogaster*, the following phylogenetic subtrees were used: ((CN, (FR, RAL)), EF), (((ZI, SD), EF), CN), and (((CN_Other, CN_LS), CN_XJ), FR). Using the subtree (((CN, AM), IS), SA) as an example, the methods for selection detection are detailed below, with similar approaches applied to other groups.

Specifically, the subtree (((CN, AM), IS), SA) was used to identify branch-specific selection signals for CN, AM, or IS, with SA as the outgroup. This analysis involved three trio tests: ((CN, AM), KN), ((CN, IS), KN), and ((AM, IS), KN). For each trio, pairwise *F*_ST_ was calculated in 4-kb sliding windows (1 kb step size) using VCFtools. PBE was subsequently calculated on the basis of established formulas^76,160^, excluding windows with fewer than 10 segregating sites. Windows in the top 1% of PBE values for each population were considered under selection.

To detect CN branch-specific selection signals, the selection signals identified for CN in the trios ((CN, AM), SA) and ((CN, IS), SA) were compared with those of other populations (AM and IS in this case). A window was considered under branch-specific selection in CN if it showed evidence of selection (top 1% PBE) in both trios for CN, but showed no selection signals (PBE values below the top 1% threshold) in the corresponding trios for the other two populations (AM and IS). Sites in these windows, with the *F*_ST_ between the CN population and the other three populations exceeding 0.1, were considered to be under selection, and the corresponding genes were identified as candidate genes. Similar methods were applied to detect selection signals for AM and IS populations. The method for selection detection in other subtrees followed the same approach described above.

### Ohana

Signals of selection at the genetic ancestry level were detected using Ohana^77,78^ version 0.1.7666.41124. Following the standard pipeline of Ohana, the autosomes and X chromosomes of *D. simulans* and *D. melanogaster* were analyzed separately. To determine the genetic ancestry proportions for each strain, the neutral sites of 681 *D. simulans* genomes and 1,335 *D. melanogaster* genomes were used. Sites and strains with a missing data rate above 10%, and sites with an MAF < 0.01 were excluded. Additionally, sites with linkage disequilibrium (LD) *r*^2^ > 0.2 within 50-kb windows were pruned by PLINK. On the basis of the ADMIXTURE results, population structure analysis was performed using qpas, with *K* = 4 for *D. simulans* and *K* = 7 and *K* = 6 for the autosomes and X chromosomes of *D. melanogaster*, respectively. The qpas parameters were set as “-mi 30 -fq” for 30 iterations, and 20 independent runs were performed to select the result with the highest log-likelihood for downstream analyses. Using the population structure results from qpas, nemeco was used to calculate the corrected allele frequencies (f-pop) for each genetic ancestry. All sites with an MAF > 0.01 were included, with nemeco parameters set to “-mi 30” for 30 iterations. These f-pop values were then used for subsequent selection analyses. To identify loci potentially under positive selection, selscan was applied with default parameters. Sites with the top 1% of log-likelihood ratio scores (LLRS) were considered candidate sites. These sites were assigned to genetic ancestry with an f-pop value exceeding 0.6, while the corresponding genes of these sites were identified as candidate genes.

### Insecticide adaptation

The list of genes associated with insecticide resistance was identical to that in a previous study^38^, which included 119 genes. Briefly, these genes were included in the GO terms “insecticide metabolic process”, “insecticide catabolic process”, and “response to insecticide” according to FlyBase, or related to insecticide resistance according to previous studies^161^. The impact of amino acid substitutions in *Ace* was estimated by ESM-scan^162^ via a web interface (https://huggingface.co/spaces/thaidaev/zsp), with the following parameters used: the “esm2_t36_3B_UR50D” model and the “higher accuracy” scoring.

### Population cage experiment with oxidative stress

To identify genetic variants associated with paraquat resistance in *D. melanogaster* and *D. simulans*, we employed a previously established approach^38^. In total, 200 and 199 isofemale strains of *D. melanogaster* and *D. simulans*, respectively, were used in this experiment. For each strain, ten 3-day-old adult males were collected and pooled into a population cage containing cornmeal food for overnight acclimation. The food was then removed, and flies were starved for 2 hours. Subsequently, two filter papers (Φ 12.5 cm), each soaked with 1.6 mL of an aqueous solution containing 5% sucrose and paraquat (50 mM for *D. melanogaster* and 20 mM for *D. simulans*), were placed into each cage. Each species was tested in two independent biological replicates. From each replicate, approximately 200 individuals with the shortest survival times (sensitive group; 205 and 153 for *D. melanogaster*, 156 and 288 for *D. simulans*) and the longest survival times (resistant group; 248 and 211 for *D. melanogaster*, 214 and 160 for *D. simulans*) were collected and subjected to whole-genome sequencing at approximately 200× per sample. In addition, two background samples per species, which is consisted of pooled DNA from 5 males per strain (approximately 1,000 individuals per species), were sequenced at approximately 400×.

Sequencing reads were aligned to the reference genomes (*D. melanogaster* dmel_r6.24 and *D. simulans*, assembled in this study) using BWA-MEM with default parameters. PCR and optical duplicates were marked using Picard. Variant calling was performed jointly across samples using FreeBayes in frequency-based pooled mode (-C 1 --pooled-continuous) with the following parameters: minimum mapping quality of 20, minimum base quality of 20, haplotype length set to 0 (-m 20 -q 20 --haplotype-length 0), maximum number of best alleles set to 4 (--use-best-n-alleles 4), and a minimum allele frequency threshold of 0.01 (-F 0.01). Variants with a quality score below 20 were discarded. Multiallelic sites were decomposed using BCFtools and vcfallelicprimitives from vcflib. SNPs located in repetitive or low-complexity regions (as described above), indels, and SNPs within 3 bp of an indel were excluded from downstream analysis. To ensure high-quality variant datasets, we applied additional filtering steps using background samples. We removed: (1) loci with sequencing depths in the top 1% (calculated separately for autosomes and the X chromosome) in background samples to avoid potential mis-mapping; (2) loci with a minor allele count (MAC) <6 for autosomes and <3 for the X chromosome in background samples; (3) loci with a MAF <0.05 in background samples; (4) loci with sequencing depth <50 in each sample; and (5) loci lacking polymorphism in at least one replicate. After filtering, 1,212,708 high-quality variants were retained in *D. melanogaster* (X: 144,303; 2L: 295,585; 2R: 229,382; 3L: 261,559; 3R: 280,757; 4: 1,122), and 2,473,893 in *D. simulans* (chrX: 265,761; chr2: 974,206; chr3: 1,231,993; chr4: 1,933).

To identify SNPs associated with paraquat resistance, we applied a Cox proportional hazards regression model using the R package survival. For each SNP, read counts of the reference (REF) and alternative (ALT) alleles were transformed into survival data, where each read was treated as an individual: REF alleles were assigned a genotype (GT) value of 0 and ALT alleles a value of 1. All reads from the sensitive group were treated as events (i.e., deaths) at the time of sampling, while reads from the resistant group were treated as censored observations. Replicates were included as separate strata in the model. The final regression formula was specified as: time ∼ GT + strata (replicate). *P* values were computed using the log-rank test and corrected for multiple testing using the Benjamini–Hochberg method separately for autosomes and the X chromosome. Following common genome-wide association study (GWAS) practices in human^163^, SNPs with an FDR < 5 × 10^-8^ were considered statistically significant in each species.

To reduce false positives arising from random associations and gene length bias, we performed a permutation test by shuffling the FDR values across loci and recalculating the number of SNPs with FDR < 5 × 10^-8^ within each gene. This procedure was repeated 1,000 times to generate a null distribution for the number of significant SNPs expected by chance for each gene. Genes in which the observed number of significant SNPs exceeded the maximum observed in the null distribution were considered to be significantly associated with oxidative stress. The convergence remained marginally significant under more stringent permutation-based criteria (*D. simulans*: 31; *D. melanogaster*: 84; overlap: 1; *P* = 0.02, hypergeometric test).

### Cross-species comparison and convergent evolution

For each gene in *D. simulans*, homology with *D. melanogaster* genes was determined through cross-validation of results from eggNOG-mapper, LiftOver, and BLASTP. A gene pair was confirmed as homologous if at least two of these methods supported the relationship, whereas both *D. melanogaster* and *D. simulans* genes exhibited a one-to-one correspondence. Using this approach, 11,827 one-to-one orthologous genes were identified between the two species. Only SNPs reside in genic regions were assigned. To assess the enrichment of overlapping target genes between the species, a hypergeometric test was used. The number of homologous genes, target genes in *D. simulans*, target genes in *D. melanogaster*, and overlapping genes between the two species were input into the R function phyper(). The parameter “lower.tail = FALSE” was specified to test whether the overlap was significantly enriched.

To assess whether the observed convergent selection signals were driven by gene flow, we compared the genomic similarity between 1,335 *D. melanogaster* and 681 *D. simulans* genomes using a *k*-mer-based approach. Reads mapped to nonrepetitive regions or the 898 convergent genes were extracted using SAMtools, and *k*-mer abundance was calculated with Sourmash version 4.8.3 (parameters: “-p k=31,abund” or “-p k=51,abund”). An angular similarity matrix for all 2,016 samples was then constructed using Sourmash with default parameters.

### Gene Ontology enrichment analysis

Gene Ontology (GO) annotations for each gene in *D. simulans* performed via using InterProScan^131^ and eggNOG-mapper^130,132^ (described before). For *D. melanogaster*, the correspondence between genes and GO terms was obtained from FlyBase (http://ftp.flybase.net/releases/current/precomputed_files/go/gene_association.fb.gz, accessed on July 6, 2023). We perform the following analysis on the basis of biological process (BP) terms only. GO enrichment analysis for candidate loci identified through selection scan (*H*_FW_, PBE and Ohana) was performed using Gowinda version 1.12. For genes under selection or associated with oxidative stress, loci under selection were used as inputs, with all polymorphic site coordinates in the corresponding species serving as the background. The analysis was restricted to the biological process category, with parameters set to “--simulations 10000000 --gene-definition gene --mode gene”. GO enrichment analysis for candidate genes of MKT or convergently adaptive genes was conducted using the enricher() function in clusterProfiler version 3.18.1, with the parameter “pAdjustMethod” set to “fdr”. The enrichment of convergently adaptive genes was performed using one-to-one orthologs of *D. simulans* as input. GO BP terms with *P* < 0.01 and at least three candidate genes were clustered in semantic space and visualized with GO-Figure! version 1.0.2 using default parameters.

### Statistical analyses

All the statistical analyses were performed using R (www.r-project.org).

## Acknowledgments

We thank the National Center for Protein Sciences at Peking University for their technical assistance. Some of the analyses were performed on the High-Performance Computing Platform of the Center for Life Sciences. The project was supported by the Ministry of Science and Technology of the People’s Republic of China grant 2022YFE0132000 (JL), Yunnan Provincial Science and Technology Project at Southwest United Graduate School grant 202302A0370006 (JL), National Natural Science Foundation of China grant 32070597 (JL) and Natural Science Foundation of Beijing grant 5212006 (JL).

## Author contributions

W.L., C.L., J.C., and Q.W. designed research. W.L., C.L., J.C., and Q.W. performed research. W.L., C.L., J.C., and Q.W. analyzed the data. W.L. and J.L. wrote the paper. All of the authors reviewed and edited the manuscript.

## Data availability statement

The sequencing data for genome assembly, population genomic, and population cage experiment have been deposited at NCBI under accessions PRJNA1244552, PRJNA1245360, and PRJNA1283574, respectively.

## Code availability statement

All software used for data analyses is publicly available. Any additional information required to reanalyze the data reported in this paper is available from the lead contact upon request.

---

**Figure S1.**
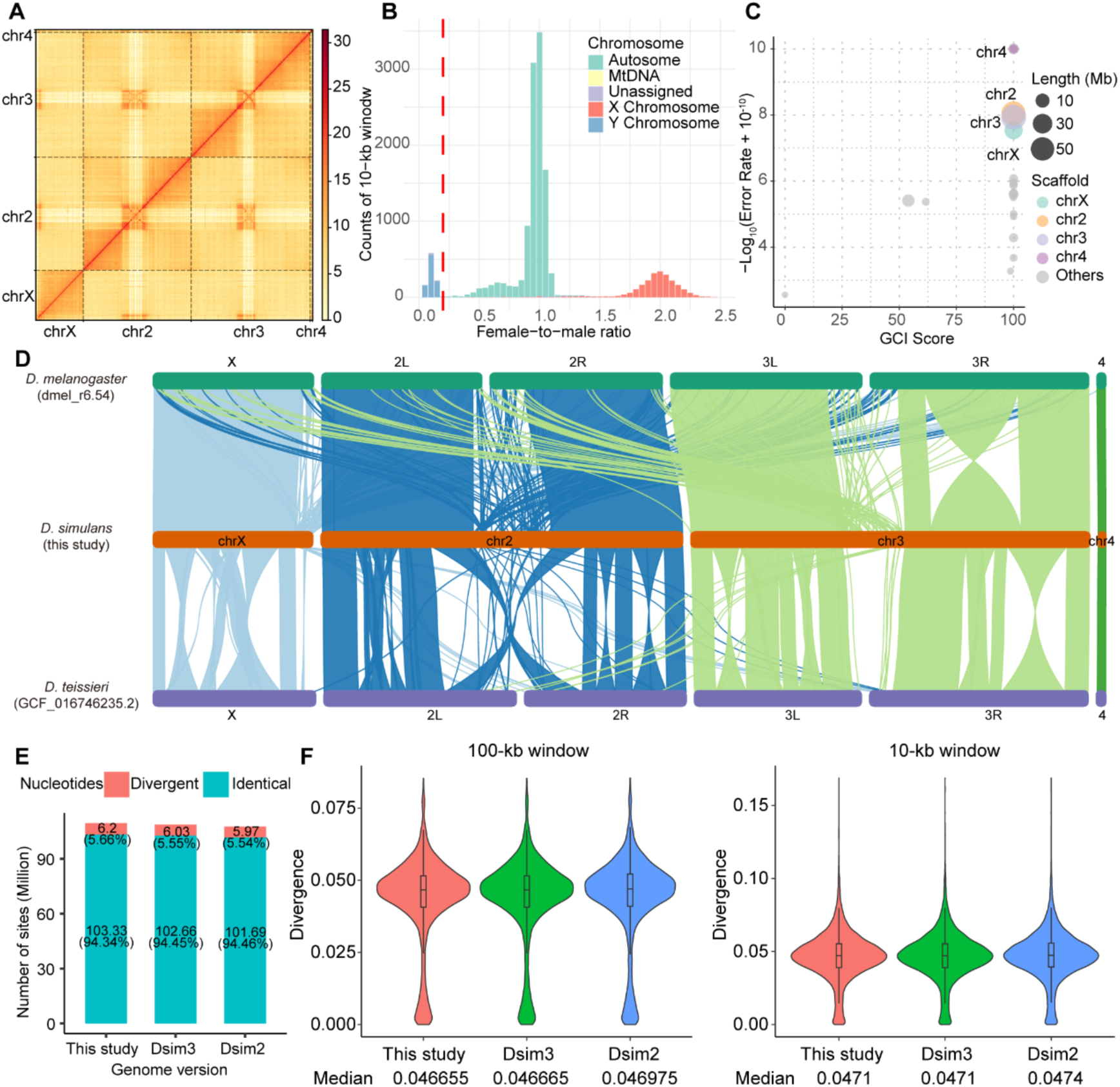
Genome quality of the genomic features of *D. simulans*. (A) High-throughput chromosome conformation capture (Hi-C) heatmap for the genome assembly of *D. simulans*. The interaction density is quantified on the basis of the number of supporting Hi-C reads and depicted using a color gradient ranging from pale yellow (low density) to dark red (high density). A heatmap of the four major chromosomes (chrX, chr2, chr3, and chr4) is shown. (B) Female‒male ratio of coverage calculated from Illumina reads. The median ratio was calculated across 10-kb windows and normalized to the total number of mapped reads to autosomes in virgin females or males. The 10-kb windows with a median ratio below 0.2 were considered Y-linked regions. (C) Assembly accuracy and continuity of the *D. simulans* genome evaluated by Merqury and the Genome Continuity Inspector (GCI) score. The size of the circle represents the length of the scaffold, while the four major chromosomes are filled in different colors. (D) Synteny analysis showing a one-to-one correspondence between the four chromosomes of the *D. melanogaster* genome r6.54 (top), the *D. simulans* genome assembly from the present study (middle), and the *D. teissieri* genome Prin_Dtei_1.1 (bottom). Each line represents an orthologous block between the compared genomes, color-coded according to their chromosome of origin. (E) Number of sites in different *D. simulans* genome assemblies that can be mapped to orthologous positions in the *D. melanogaster* genome. For each assembly, the number and proportion of sites with identical or divergent nucleotides at orthologous positions are indicated. (F) Divergence between the *D. melanogaster* (Dmel6) and *D. simulans* genomes assembled in this study and from previous studies (Dsim3 and Dsim2), as calculated using 100-kb and 10-kb nonoverlapping sliding windows. The levels of divergence from Dmel6 were comparable across the different *D. simulans* genome assemblies. In the violin plots, the boxes represent the interquartile range (25th to 75th percentiles), the horizontal lines within the boxes indicate the median, and the whiskers denote the minimum and maximum values.

**Figure S2.**
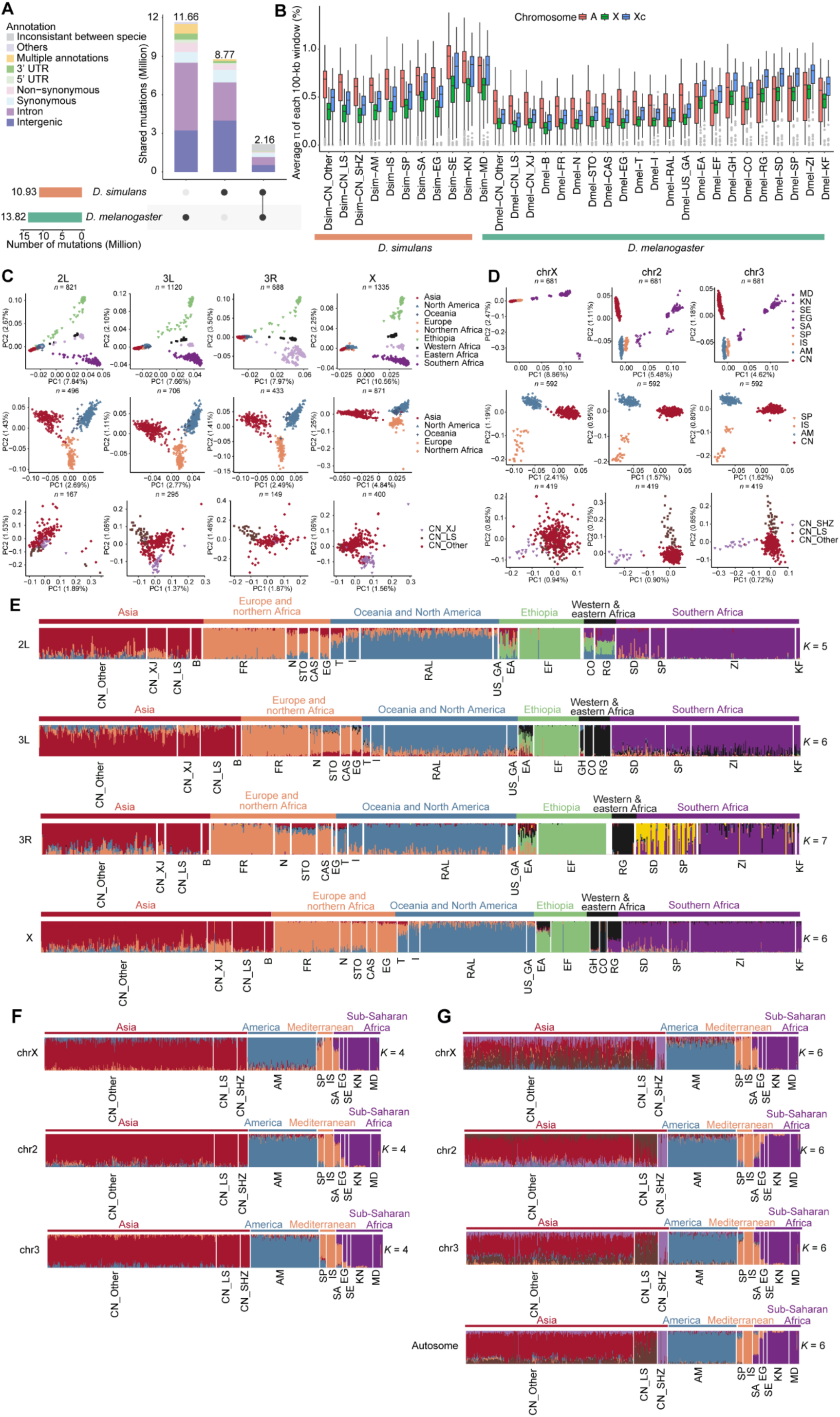
Genetic diversity and population stratification of *D. melanogaster* and *D. simulans*. (A) Overlap of SNPs identified in *D. simulans* and *D. melanogaster*. Stacked bar graphs showing the number of SNPs of different annotation types. (B) Genetic diversity (π) of 11 populations of *D. simulans* and 22 populations of *D. melanogaster*. The genetic diversity of autosomes (π_A_), the X chromosome (π_X_) and the X chromosome after correction for population size (π_Xc_) were calculated on the basis of 200-kb nonoverlapping windows using all SNPs for each population. The genetic diversity of all autosomes was combined as π_A_. (**C** and **D**) Principal component analysis (PCA) results on the basis of neutral sites from inversion-free *D. melanogaster* strains on each chromosomal arm (**C**), and from *D. simulans* strains on each chromosome (**D**). The number of genomes (*n*) is labeled on the graph. The proportion of variance explained by each corresponding principal component is indicated in parentheses. The first two PCs were plotted for global strains (upper), cosmopolitan strains (middle), and CN strains (bottom). (E) The ancestry compositions of each strain of *D. melanogaster* with the best-fitting *K* for chromosomes 2L, 3L, 3R, and X. The results were based on neutral sites from inversion-free chromosome arms. The optimal number of ancestries (*K*) was determined when the minimum cross validation (CV) error median was reached (see also Table S13). (**F** and **G**) The ancestry compositions of each strain of *D. simulans* with *K* = 4 (**F**) and *K* = 6 (**G**) for chr2, chr3, and chrX. The results were based on neutral sites. Additional genetic ancestries were found in the CN_SHZ and CN_LS populations when *K* = 6.

**Figure S3.**
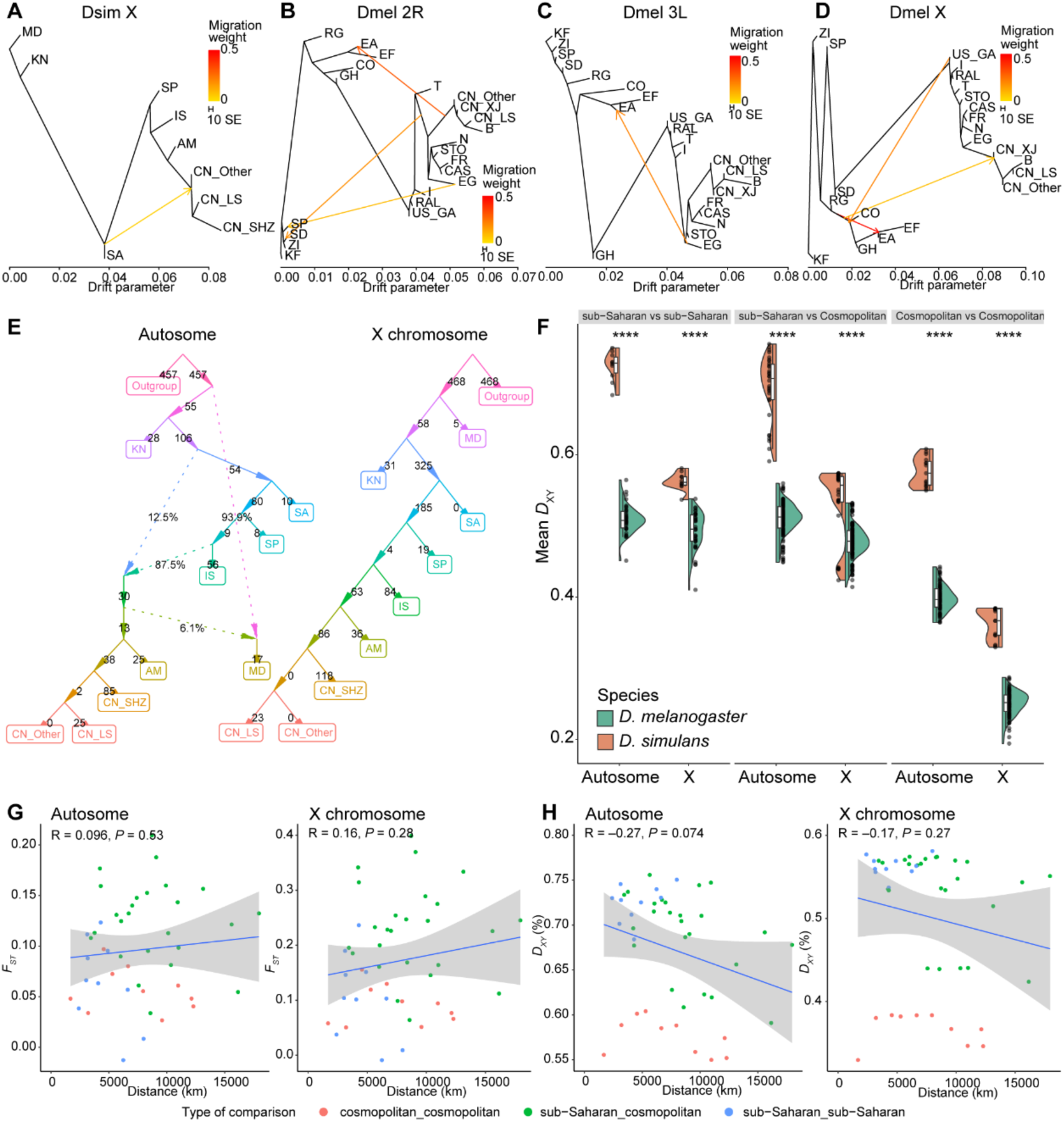
Gene flow and population differentiation of *D. melanogaster* and *D. simulans*. (**A**–**D**) Maximum likelihood tree and gene flow inferred using TreeMix. The optimal number of migration edges (*m*) was determined by OptM. (A) Results based on neutral sites of the X chromosome from nine populations of *D. simulans* with population sizes larger than 10. The migration edge was inferred from the common ancestor of SA and cosmopolitan populations to the ancestor of CN. (B) Results based on neutral sites of inversion-free 2R from 22 populations of *D. melanogaster*. Three migration edges were inferred: 1) from the common ancestor of Asian populations to EA; 2) from the common ancestor of Asian and European populations to SD; and 3) from EG to SP. (C) Results based on neutral sites of inversion-free 3L from 22 populations of *D. melanogaster*. The migration edge was inferred from the ancestor of EG to EA. (D) Results based on neutral sites of the X chromosome from 22 populations of *D. melanogaster*. Three migration edges were inferred: 1) from the common ancestor of CO, GH, EA and EF to the common ancestor of Asian populations; 2) from the common ancestor of CO, GH, EA and EF to the common ancestor of Ethiopian populations; and 3) from US_GA to the common ancestor of CO, GH, EA and EF. (E) Admixture graph of nine populations of *D. simulans* with population sizes larger than 10, estimated by ADMIXTOOLS2 according to the likelihood score based on neutral SNPs of the autosomes and X chromosome. The population split is represented by solid lines, and branch lengths are provided in units of 10 × *f*2 drift distance. The inferred admixture events are shown by dotted lines with proportions annotated. (F) Split violin plots exhibiting genetic differentiation of *D. simulans* (orange) and *D. melanogaster* (green) measured as *D*_XY._ The *D*_XY_ was calculated on the basis of all sites from all strains for autosomes or the X chromosome. The top and bottom of the boxes within the split violin depict the 75^th^ and 25^th^ percentiles of the distribution, respectively. The horizontal lines within the boxes signify the median values. Axis endpoints are labeled by the minimum and maximum values. The statistical significance according to the Wilcoxon rank sum exact test is shown for each comparison. ns *P* > 0.05, *** *P* < 0.001, and **** *P* < 0.0001. (**G** and **H**) The correlation between geographic distance and genetic differentiation of *D. simulans* measured by *F*_ST_ (**G**) and *D*_XY_ (**H**) on autosomes and the X chromosome, respectively. The coefficient R and *P* value of the linear regression using the Pearson correlation method are shown.

**Figure S4.**
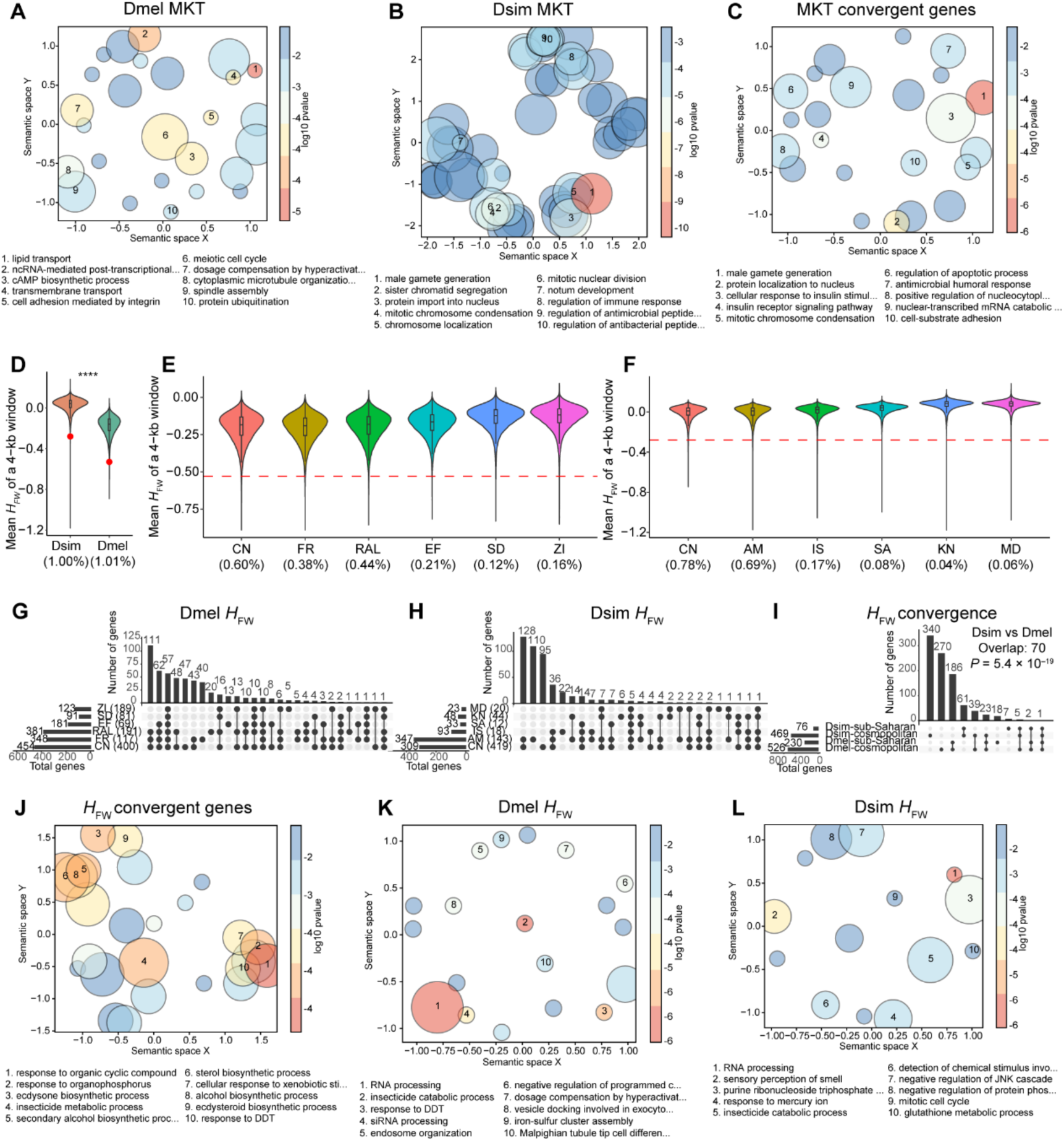
Natural selection detected by the McDonald–Kreitman test (MKT) and Fay and Wu’s *H* (*H*_FW_) in *D. melanogaster* and *D. simulans*. (A) Scatterplot summarizing 44 Gene Ontology biological process (GO BP) terms enriched in 620 adaptive genes detected by MKT in *D. melanogaster*. A description of the top 10 representative GO terms is provided below the graph, and the detailed results are provided in Table S18. Semantically similar GO terms are positioned closer together in the plot. Bubble size and color represent the number of grouped GO terms and log-transformed *P* values, respectively. (B) Scatterplot summarizing 154 GO BP terms enriched in 1255 adaptive genes detected by MKT in *D. simulans*. (C) Scatterplot summarizing 70 GO BP terms enriched in 208 convergent genes detected by MKT in the two species. (D) Distribution of the mean *H*_FW_ of 4-kb windows in six major populations of *D. simulans* and *D. melanogaster*. Compared with *D. melanogaster*, *D. simulans* has a significantly greater *H*_FW_. The red dots represent the thresholds (*D. simulans*: *H*_FW_ < −0.28; *D. melanogaster*: *H*_FW_ < −0.53) used to determine whether a window was under natural selection. The percentages below the species names represent the proportion of windows under the specific threshold. The calculation of *H*_FW_ includes all the SNPs and all the strains of the six major populations. *P* values were derived from Wilcoxon tests, **** *P* < 0.0001. (**E** and **F**) Distribution of the mean *H*_FW_ of 4-kb windows (step 1 kb) in six major populations of *D. melanogaster* (**E**) and *D. simulans* (**F**). The 4-kb windows with fewer than 50 segregating sites were discarded. The dashed line in red represents the threshold (*D. simulans*: *H*_FW_ < −0.28; *D. melanogaster*: *H*_FW_ < −0.53) used to determine whether a window was under natural selection. (**G** and **H**) Overlap of positively selected genes detected by *H*_FW_ in the six major populations of *D. melanogaster* (**G**) and *D. simulans* (**H**). The number of genomes used in the analysis for each population is indicated in parentheses. (I) Overlap of positively selected genes detected by *H*_FW_ in *D. simulans* and *D. melanogaster*. Populations were grouped according to their geographic locations of sampling in each species, and each group included three populations. The statistical significance according to the hypergeometric test is shown. _(J)_ Scatterplot summarizing 54 GO BP terms enriched in 70 convergent genes detected by *H*_FW_ in the two species. _(K)_ Scatterplot summarizing 23 GO BP terms enriched in 544 adaptive genes detected by *H*_FW_ in *D. melanogaster*. (L) Scatterplot summarizing 45 GO BP terms enriched in 478 adaptive genes detected by *H*_FW_ in *D. simulans*. The GO enrichment analysis in panels **A**–**C** and **J** was conducted using clusterProfiler, and that in panels **K** and **L** was conducted using Gowinda.

**Figure S5.**
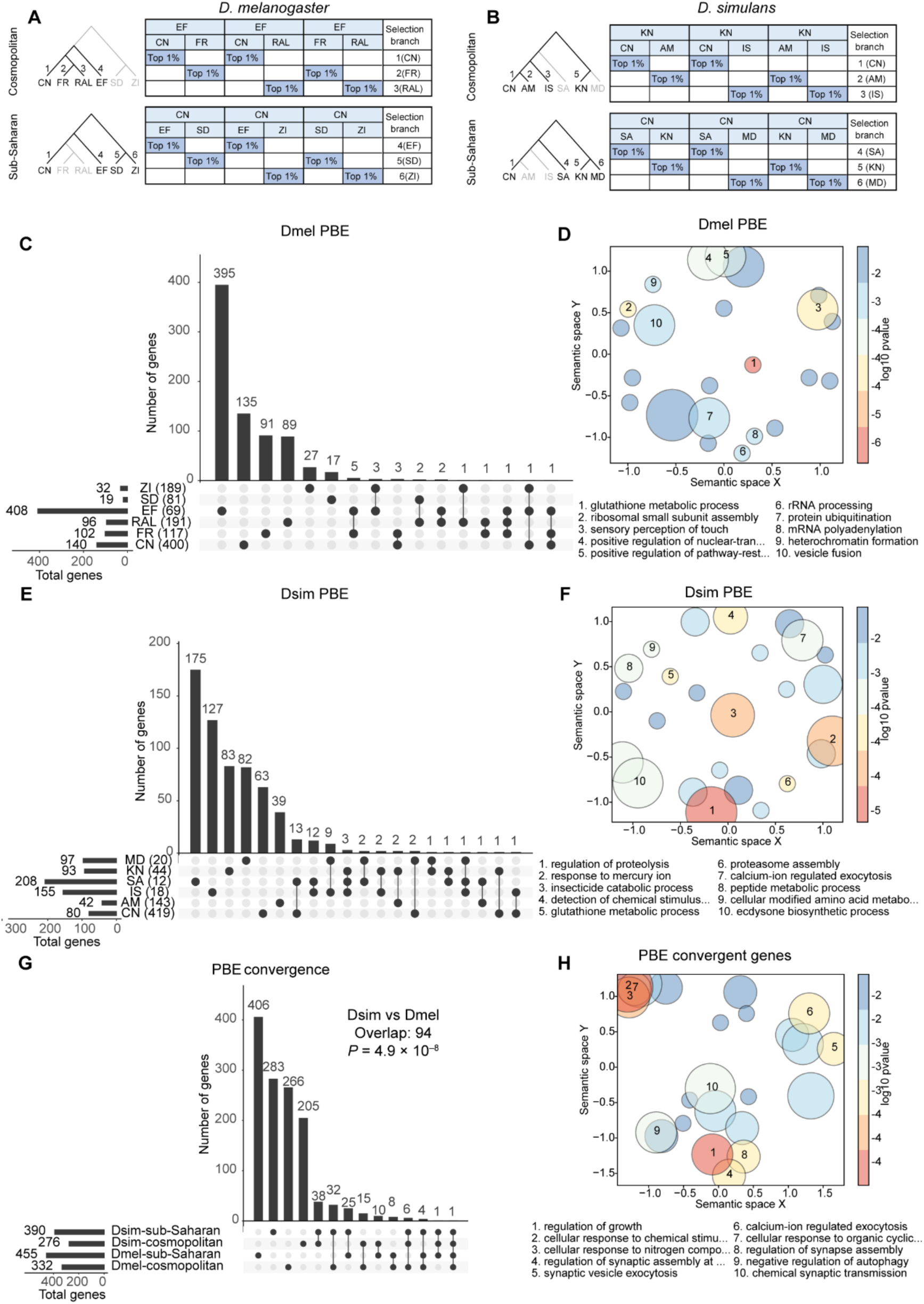
Natural selection detected by population branch excess (PBE) in *D. melanogaster* and *D. simulans*. (**A** and **B**) The selection scan procedure and the phylogenetic tree used for PBE analysis in *D. melanogaster* (**A**) and *D. simulans* (**B**). (C) Overlap of positively selected genes detected by PBE in the six major populations of *D. melanogaster.* The number of genomes used in the analysis for each population is indicated in parentheses. (D) Scatterplot summarizing 31 Gene Ontology biological process (GO BP) terms enriched in 1045 adaptive genes detected by PBE in *D. melanogaster*. A description of the top 10 representative GO terms is provided below the graph, and the detailed results are provided in Table S18. Semantically similar GO terms are positioned closer together in the plot. Bubble size and color represent the number of grouped GO terms and log-transformed *P* values, respectively. (E) Overlap of positively selected genes detected by PBE in the six major populations of *D. simulans*. (F) Scatterplot summarizing 78 GO BP terms enriched in 620 adaptive genes detected by PBE in *D. simulans*. (G) Overlap of positively selected genes detected by PBE in *D. simulans* and *D. melanogaster*. Populations were grouped according to their geographic locations of sampling in each species, and each group included three populations. The statistical significance according to the hypergeometric test is shown. (H) Scatterplot summarizing 59 GO BP terms enriched in 94 convergent genes detected by PBE in the two species. The GO enrichment analysis in panels **D** and **F** was conducted using Gowinda, and that in panel **H** was conducted using clusterProfiler.

**Figure S6.**
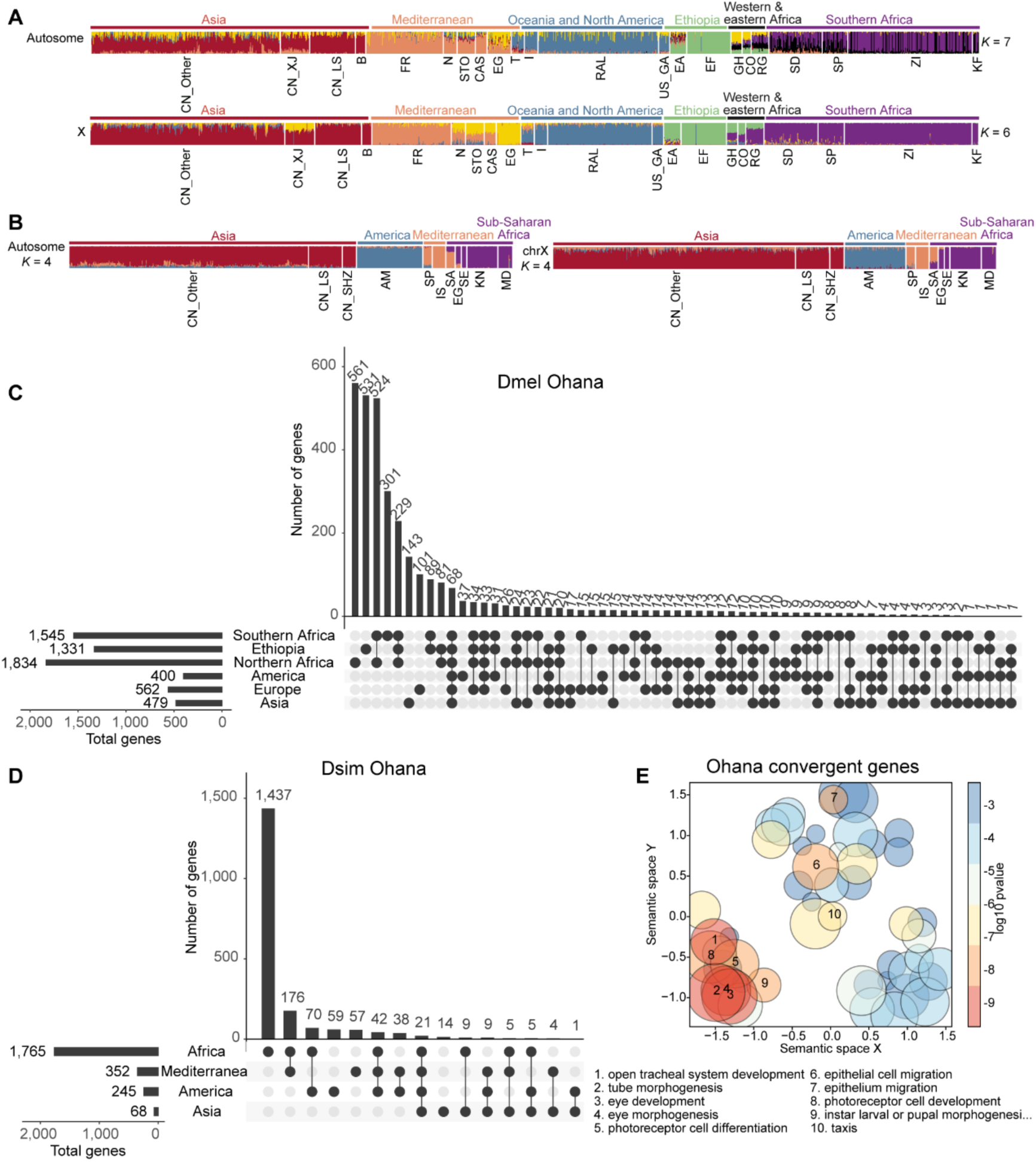
Natural selection detected by Ohana in *D. melanogaster* and *D. simulans*. (**A** and **B**) Ancestry admixture inference of *D. melanogaster* (**A**) and *D. simulans* (**B**) by Ohana. Ohana was conducted for autosomes and the X chromosome separately. The ancestry compositions of each strain are shown. The optimal number of ancestries, *K*, is determined according to the ADMIXTURE analysis. The structure is based on neutral SNPs with a MAF greater than 0.01. The linkage disequilibrium was pruned in 50-kb windows with a threshold of *r*^2^ < 0.2 by PLINK. Samples with a missing data rate that exceeded 0.1 were removed. (**C**) Overlap of selected genes detected by Ohana in the six genetic ancestries of *D. melanogaster*. SNPs whose log-likelihood ratio score (LLRS) was the highest at 1% were considered under positive selection, and genes harboring these SNPs were identified as being under positive selection. (**D**) Overlap of selected genes detected by Ohana in the four genetic ancestries of *D. simulans*. (**E**) Scatterplot summarizing 611 GO BP terms enriched in 284 convergent genes detected by Ohana in the two species. A description of the top 10 representative GO terms is provided below the graph, and the detailed results are provided in Table S18. Semantically similar GO terms are positioned closer together in the plot. Bubble size and color represent the number of grouped GO terms and log-transformed *P* values, respectively. The GO enrichment analysis in panel E was conducted using clusterProfiler.

**Figure S7.**
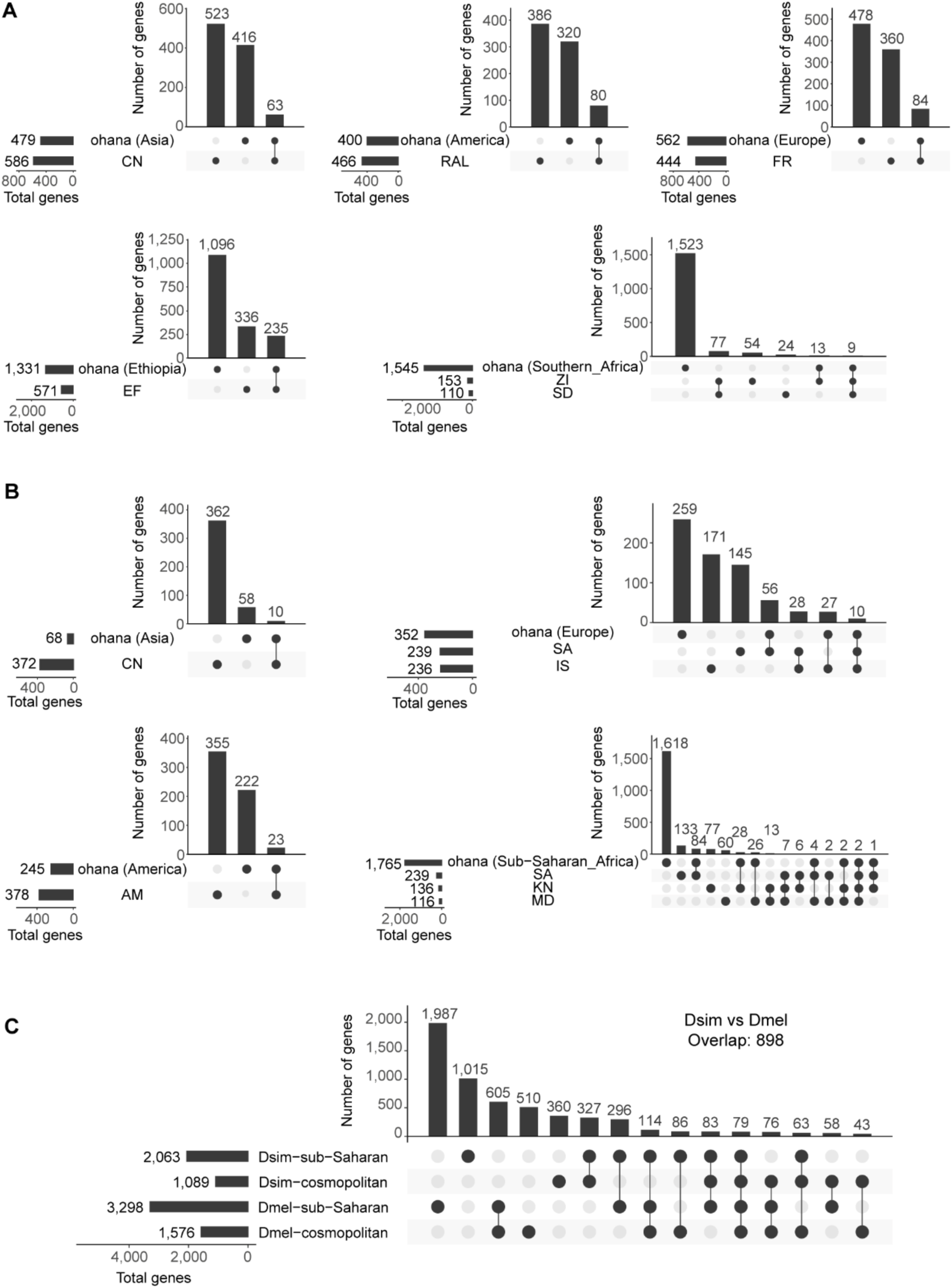
Overlap of selection signals detected by different methods within and between species. (**A** and **B**) Overlap of positively selected genes detected by Ohana in genetic ancestry and genes detected by *H*_FW_ and PBE in corresponding populations in *D. melanogaster* (**A**) and *D. simulans* (**B**). (**C**) Overlap of positively selected genes detected by *H*_FW_, PBE and Ohana in *D. simulans* and *D. melanogaster*. Populations or genetic ancestries were grouped according to their geographic locations of sampling in each species.

**Figure S8.**
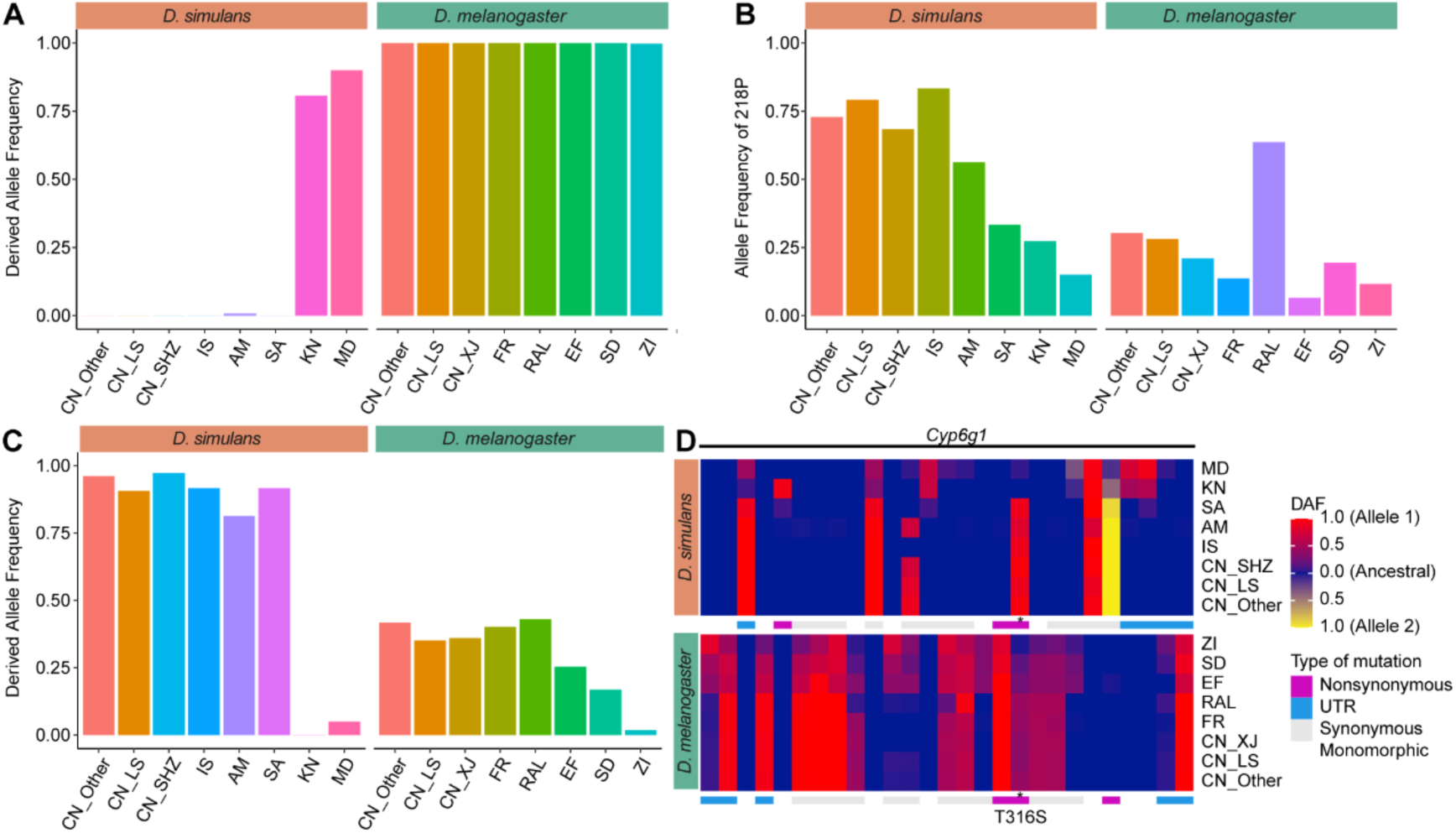
Molecular parallel adaptation in *pcm*, *CG6933*, and *Cyp6g1*. (A) Derived allele frequencies of A1513T in *D. simulans* and A1515T in *D. melanogaster* in *pcm*. The allele frequencies of six major populations of each species are shown, with three subpopulations from China shown separately. The allele in the most recent common ancestor (MRCA) of *D. simulans* and *D. melanogaster* was considered the ancestral allele. (B) Allele frequencies of 216P in *CG6933* in populations of *D. simulans* and *D. melanogaster*. The allele in the MRCA of *D. simulans* and *D. melanogaster* could not be determined. (C) Derived allele frequencies of T316S in *Cyp6g1* in populations of *D. simulans* and *D. melanogaster*. (D) The pattern of derived allele frequencies of *Cyp6g1* in populations of *D. simulans* (upper panel) and *D. melanogaster* (lower panel). Each column is an exonic SNP with minor allele frequencies over 0.05 in at least one species (calculated with all strains in each species), and each row is a population or a subpopulation. The allele in the MRCA of *D. simulans* and *D. melanogaster* was considered the ancestral allele, while the sites without ancestral information were removed. The color gradient represents the derived allele frequency of each locus in each population, with red and yellow representing two different alleles in *D. simulans* and *D. melanogaster*. The annotation of each mutation in each species is shown below the allele frequency heatmap in each panel. The asterisk denotes the missense variants that were under selection in both species.

**Figure S9.**
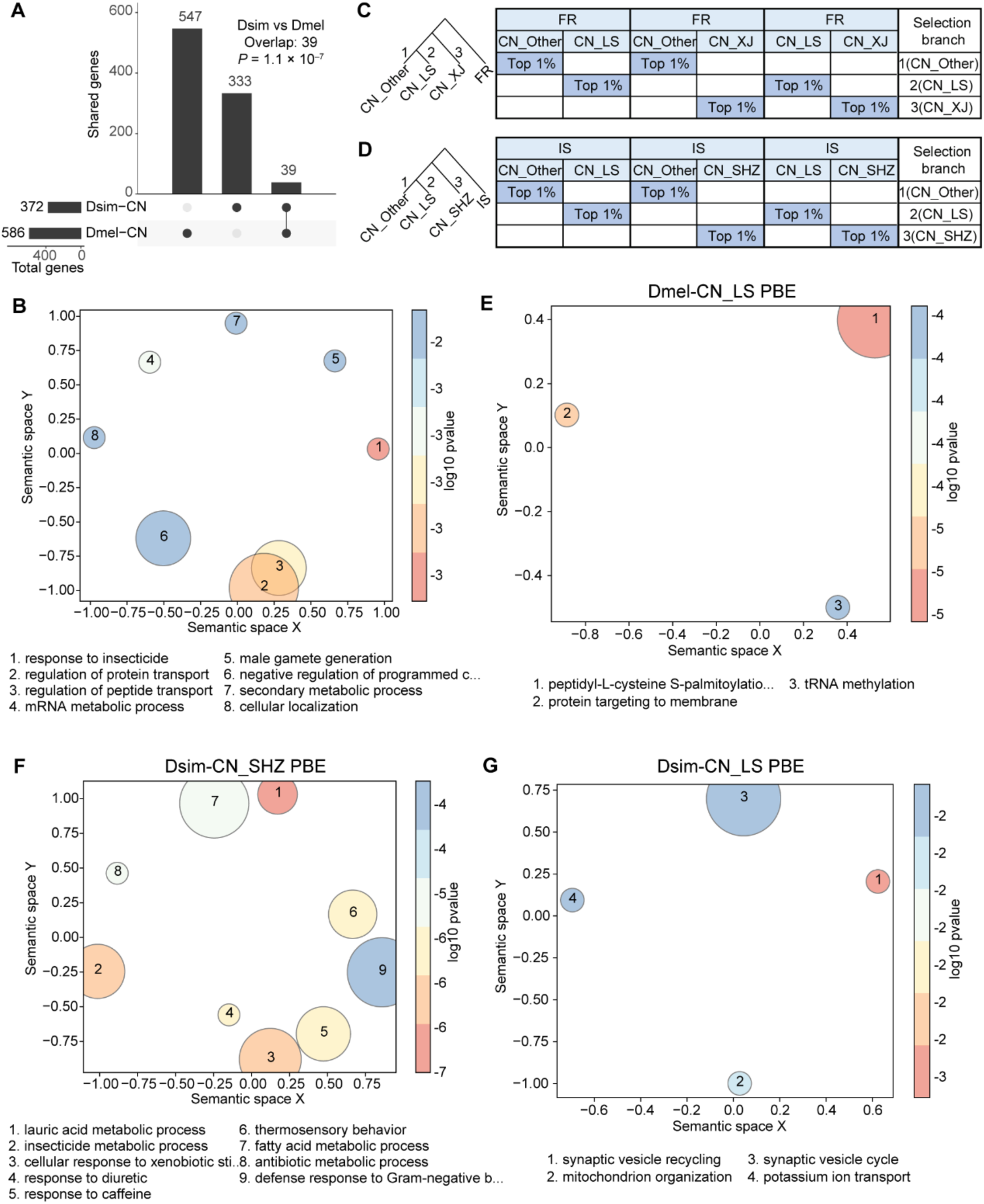
Natural selection in populations from China in *D. melanogaster* and *D. simulans*. (A) Overlap of positively selected genes detected by Fay and Wu’s *H* (*H*_FW_) and population branch excess (PBE) methods in the CN populations of *D. simulans* and *D. melanogaster*. The statistical significance of the hypergeometric test is shown. (B) Scatterplot summarizing 12 Gene Ontology biological process (GO BP) terms enriched in 39 adaptive genes detected by PBE in the CN population of two species. A description of the top representative 10 GO terms is provided below the graph, and the detailed results are provided in Table S18. Semantically similar GO terms are positioned closer together in the plot. Bubble size and color represent the number of grouped GO terms and log-transformed *P* values, respectively. (**C** and **D**) The selection scan procedure and the phylogenetic tree used for PBE analysis in the CN subpopulations of *D. melanogaster* (**C**) and *D. simulans* (**D**). (E) Scatterplot summarizing four GO BP terms enriched in 204 adaptive genes detected by PBE in CN_LS of *D. melanogaster* (Dmel-CN_LS). (F) Scatterplot summarizing 41 GO BP terms enriched in 191 adaptive genes detected by PBE in CN_SHZ of *D. simulans* (Dsim-CN_SHZ). (G) Scatterplot summarizing five GO BP terms enriched in 55 adaptive genes detected by PBE in CN_SHZ of *D. simulans* (Dsim-CN_SHZ). The GO enrichment analysis in panels **E**–**G** was conducted using Gowinda, and that in panel B was conducted using clusterProfiler.

**Figure S10.**
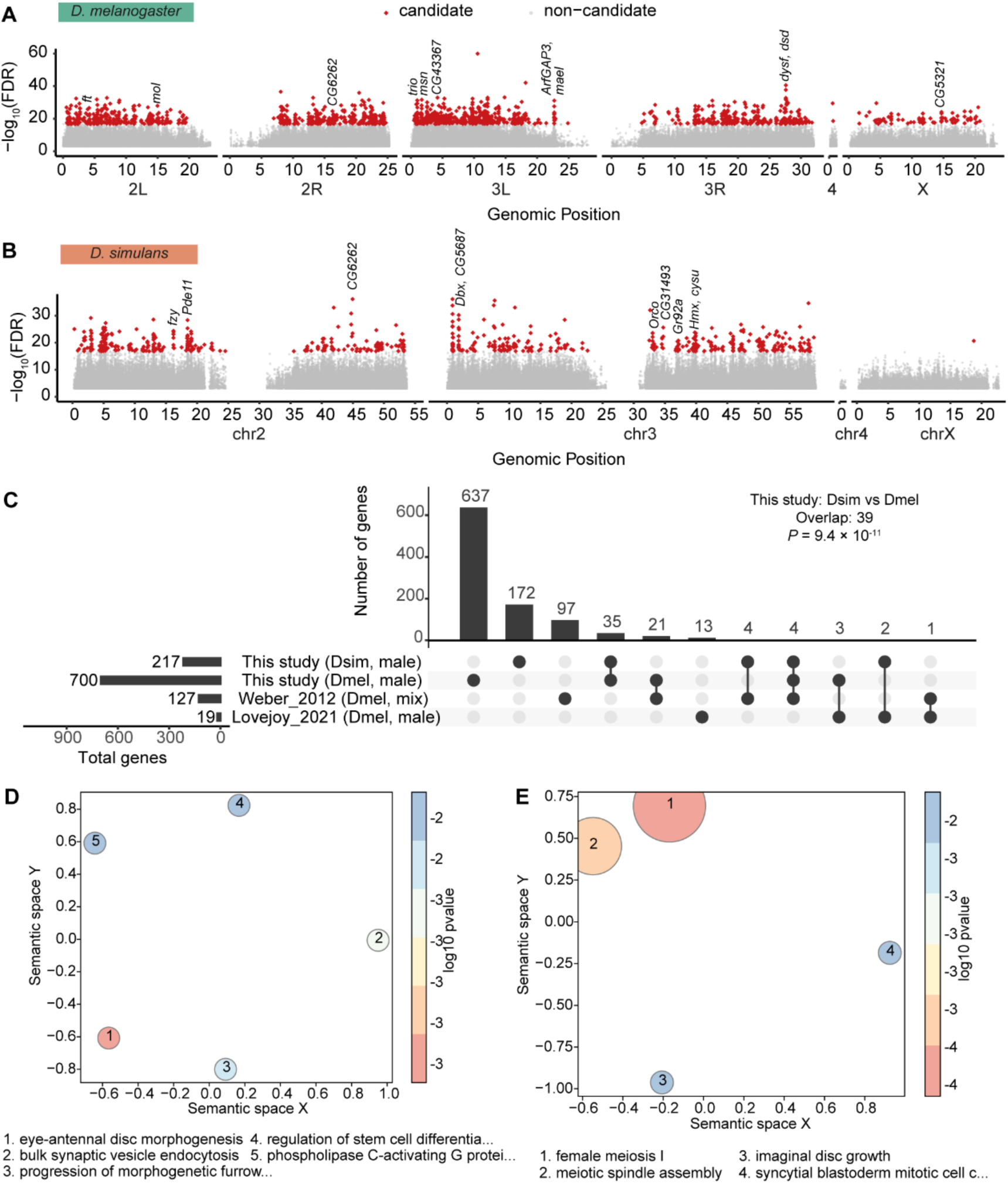
Convergent genomic response to oxidative stress. (**A** and **B**) Manhattan plot of candidate SNPs associated with oxidative stress in *D. melanogaster* (**A**) and *D. simulans* (**B**). *P* values were calculated by the Cox Proportional Hazards regression model, while Multiple testing correction was applied using the Benjamini-Hochberg method to calculate FDR. The candidate SNPs (FDR < 5 × 10^-8^) were colored in red, and only SNPs with FDR < 0.05 were shown. (C) Overlap of associated genes between *D. simulans*, *D. melanogaster*, and previous studies^86,87^. The species and sex of flies is indicated in parentheses. The statistical significance based on the hypergeometric test is shown. (**D** and **E**) Scatterplot summarizing 12 Gene Ontology biological process (GO BP) terms enriched in 700 and 217 genes associated with oxidative stress in *D. melanogaster* (**D**) and *D. simulans* (**E**). A description of detailed results is provided in Table S18. Semantically similar GO terms are positioned closer together in the plot. Bubble size and color represent the number of grouped GO terms and log-transformed *P* values, respectively. The GO enrichment analysis was conducted using Gowinda.

**Figure S11.**
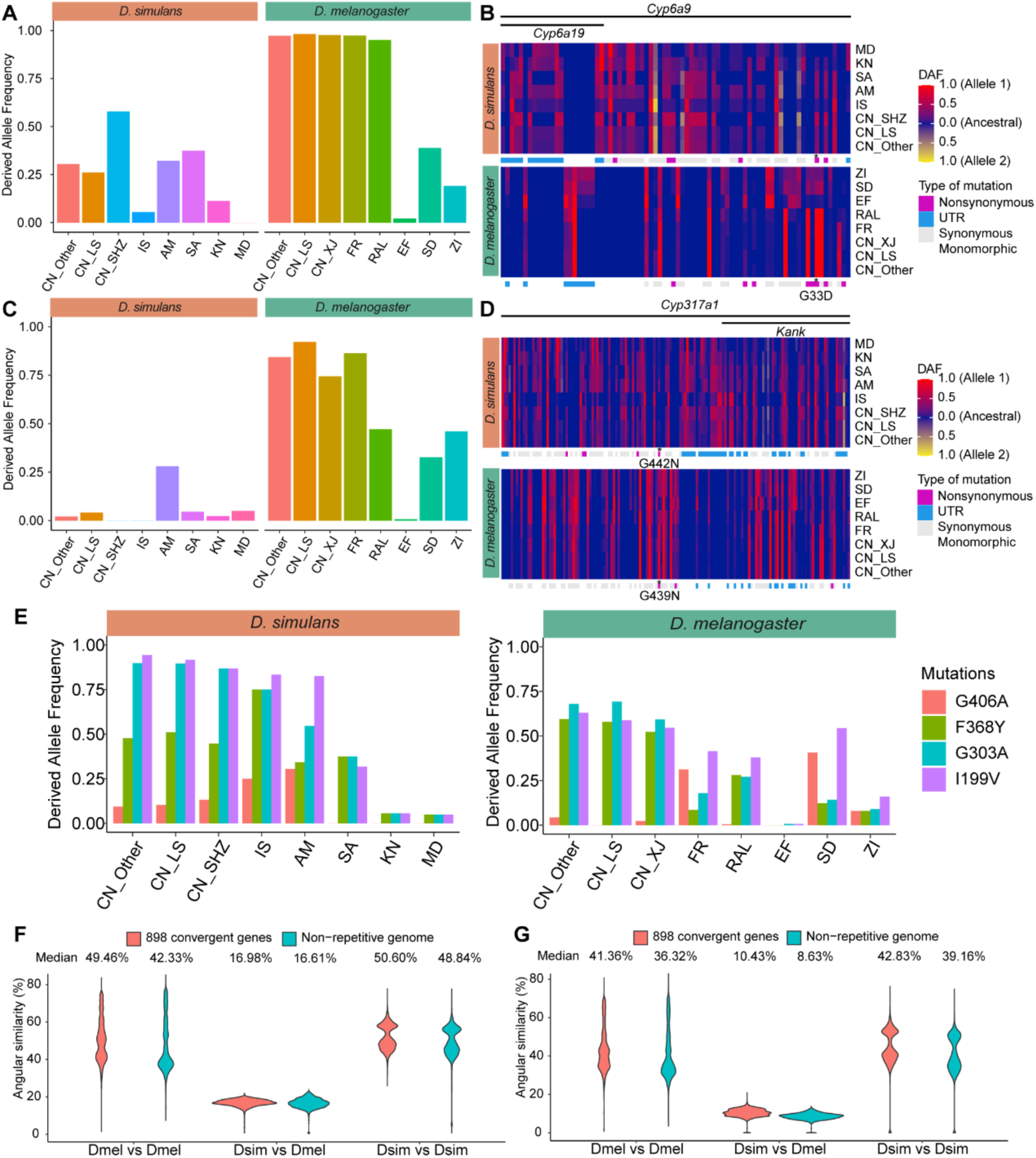
Molecular parallel adaptation in insecticide resistance genes. (A) Derived allele frequencies of G33D in *Cyp6a9* in populations of *D. simulans* and *D. melanogaster*. (B) The pattern of derived allele frequencies of *Cyp6a9* in populations of *D. simulans* (upper panel) and *D. melanogaster* (lower panel). Each column is an exonic SNP with minor allele frequencies over 0.05 in at least one species (calculated with all strains in each species), and each row is a population or a subpopulation. The allele in the MRCA of *D. simulans* and *D. melanogaster* was considered the ancestral allele, while the sites without ancestral information were removed. The color gradient represents the derived allele frequency of each locus in each population, with red and yellow representing two different alleles in *D. simulans* and *D. melanogaster*. The annotation of each mutation in each species is shown below the allele frequency heatmap in each panel. The asterisk denotes the missense variants that were found in both species. (C) Derived allele frequencies of G442N in *D. simulans* and G439N in *D. melanogaster* in *Cyp317a1*. (D) The pattern of derived allele frequencies of *Cyp317a1* in populations of *D. simulans* and *D. melanogaster*. (E) Derived allele frequencies of I119V, G303A, F368Y and G406A of *Ace* in populations of *D. simulans* and *D. melanogaster*. (**F** and **G**) Violin plots comparing angular *k*-mer similarity among 2,016 *D. melanogaster* and *D. simulans* samples using *k* = 31 (**F**) and *k* = 51 (**G**). Angular similarities were consistently greater in within-species comparisons (“Dmel vs. Dmel” and “Dsim vs. Dsim”) than in interspecific comparisons (“Dmel vs. Dsim”), both in nonrepetitive genomic regions and across the 898 convergent genes.

**Figure S12.**
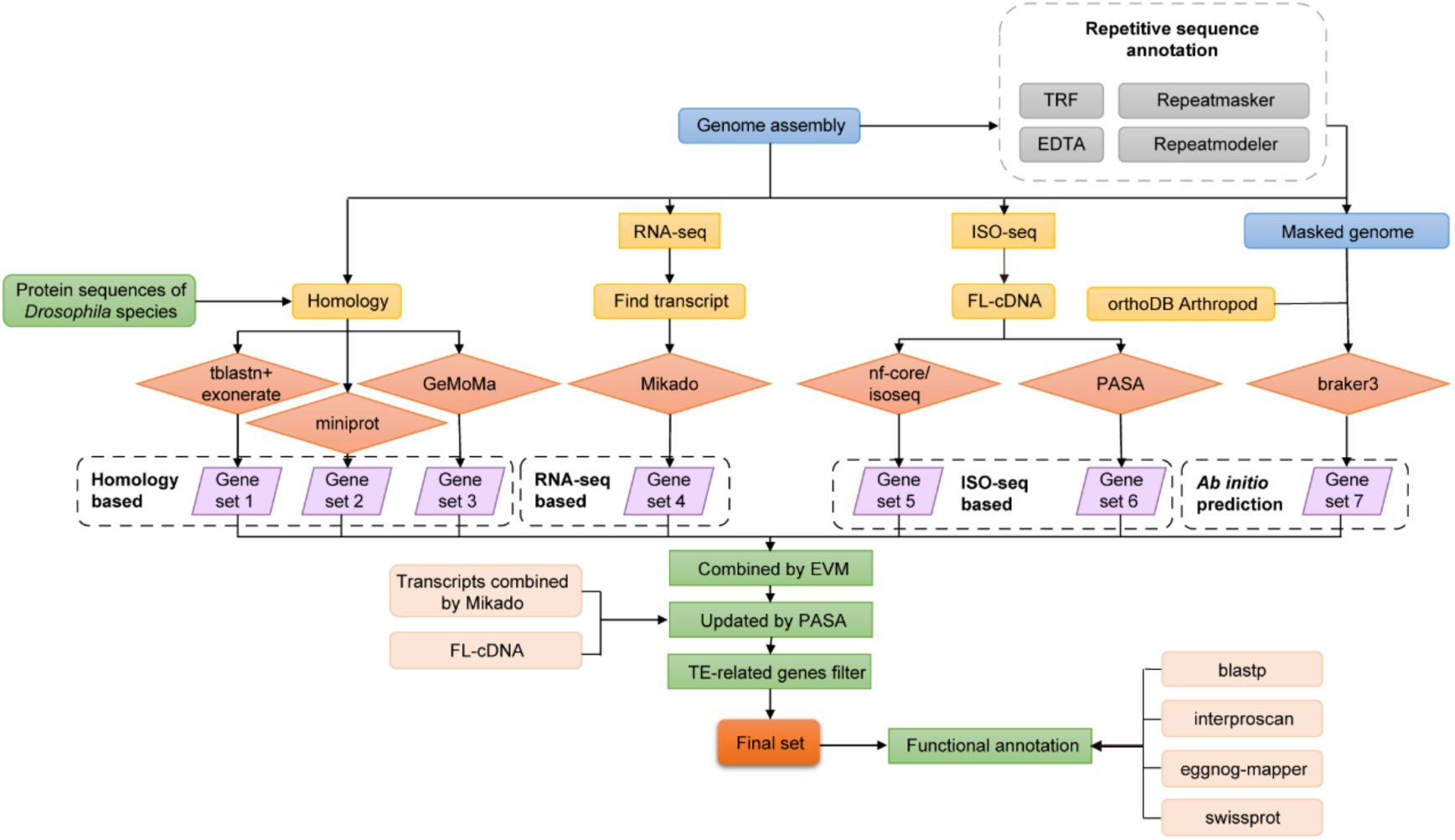
Flowchart of the method for protein-coding gene annotation in the *D. simulans* genome. Four different approaches were used to generate seven gene sets, including protein-based homology search (gene sets 1–3), RNA sequencing (RNA-seq)-based prediction (gene set 4), ISO-seq-based prediction (gene sets 5 and 6), and *ab initio* prediction (gene set 7). The seven gene sets were further combined using Evidence Modeler, and functional annotation was performed.

**Figure S13.**
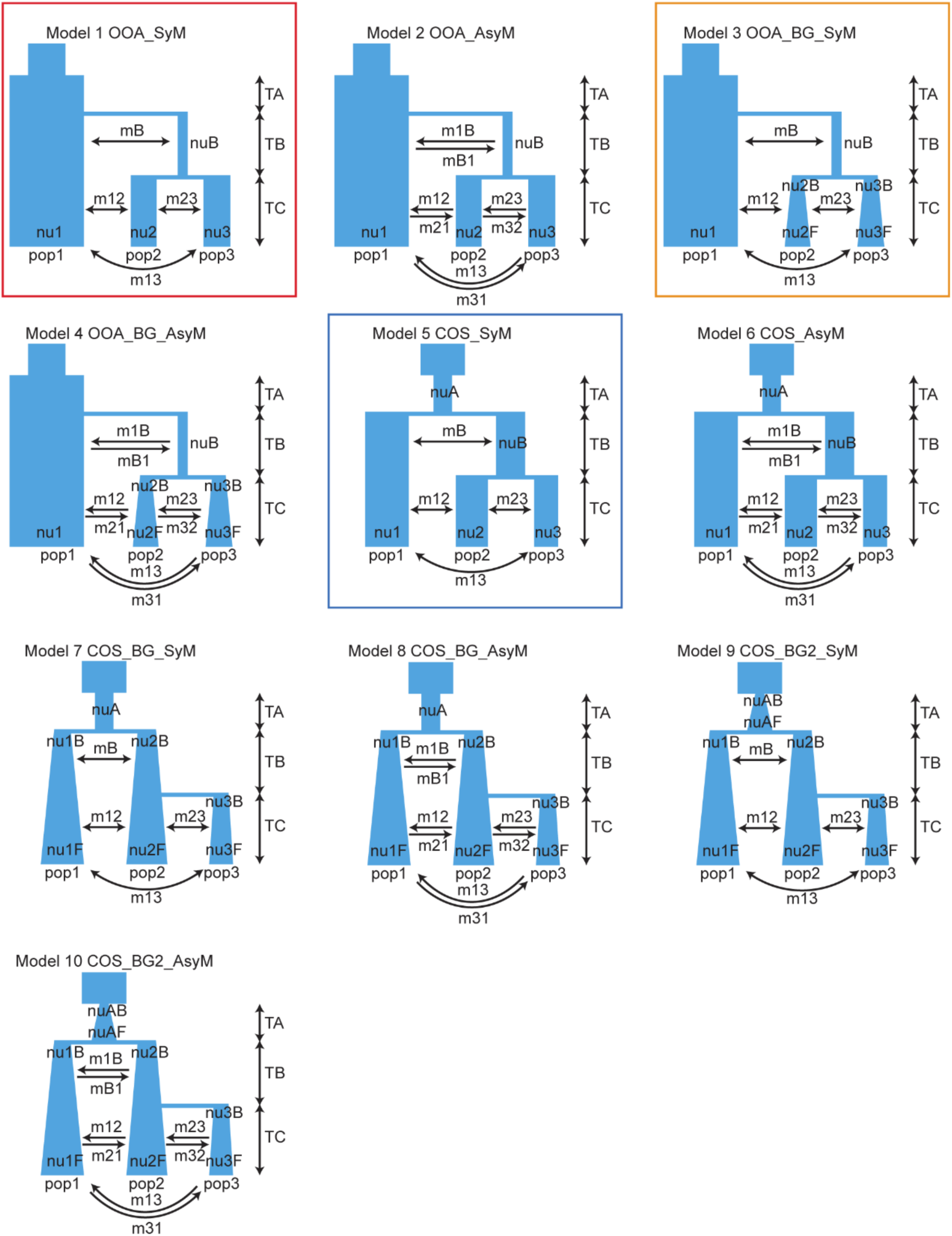
Schematics of the demographic models tested using moments. For ((CN_LS, IS), MD), ((AM, IS), MD), and ((SP, IS), MD), the best model chosen from models 1–4 is model 1, indicated by the red box. For ((IS, KN), MD), the best model chosen from models 1– 4 is model 3, indicated by the orange box. For ((CN_LS, CN_Other), IS) and ((CN_SHZ, CN_Other), IS), the best model chosen from models 5–10 is model 5, indicated by the blue box. All parameters of the model are shown in schematics, including the population size at each time point (nu), the migration rate between populations (m), and the duration of each event (T).

